# Molecular basis of egg coat cross-linking sheds light on ZP1-associated female infertility

**DOI:** 10.1101/591966

**Authors:** Kaoru Nishimura, Elisa Dioguardi, Shunsuke Nishio, Alessandra Villa, Ling Han, Tsukasa Matsuda, Luca Jovine

## Abstract

Interaction between sperm and the egg zona pellucida (ZP) is the first step of mammalian fertilization, and ZP component ZP1 is important for fertility by covalently cross-linking ZP filaments into a matrix. Like ZP4, a structurally-related subunit absent in the mouse, ZP1 is predicted to contain an N-terminal ZP-N domain of unknown function. Characterization of ZP1 proteins carrying mutations from infertile patients suggests that, unlike in the mouse, filament cross-linking by ZP1 is crucial for human ZP assembly. We map the function of ZP1 to its ZP-N1 domain and determine crystal structures of ZP-N1 homodimers from a chicken homolog of ZP1. These reveal that ZP filament cross-linking is highly plastic and can be modulated by ZP1 fucosylation and, potentially, zinc sparks. Moreover, we show that ZP4 ZP-N1 forms non-covalent homodimers in chicken but not human. Together, these data identify human ZP1 cross-links as a promising target for non-hormonal contraception.

## Introduction

Vertebrate oocytes are surrounded by a specialized extracellular coat that is referred to as zona pellucida in mammals (ZP) and vitelline envelope in non-mammals (VE). This matrix plays essential roles in oogenesis and provides a physical barrier that protects the developing embryo; in tetrapods, it also mediates the initial interaction between gametes at fertilization and, in mammals, contributes to the block to polyspermy^1,2^. Although their fine morphological appearance differs between classes, all vertebrate egg coats consist of filaments made up of a variable number of glycoprotein components that polymerize using a common ZP module^3,4^. This unit, which consists of two structurally related immunoglobulin (Ig)-like domains (ZP-N and ZP-C), is also the building block of the generally more complex coats of invertebrate eggs; moreover, it is contained in many additional extracellular proteins not involved in fertilization^5^.

The human ZP contains four glycoproteins (hZP1-4), only three of which (mZP1-3) are found in the mouse due to pseudogenization of *Zp4*^6–8^. While mZP2 and mZP3 are the major components of ZP filaments^6^ and have long been implicated in the interaction with sperm^9–11^, less abundant mZP1 was shown to covalently cross-link ZP filaments by forming homodimers held together by intermolecular disulphide bridge(s)^12^. This architecture is consistent with the observation that *Zp2* or *Zp3* null mice lack a ZP^13–15^, whereas *Zp1* null animals have a fragile and loosen egg coat^16^. Moreover, the structural similarity between the ZP modules of ZP2 and ZP1^5,17^ suggests that cross-link points are introduced into ZP filaments when mZP1 is occasionally incorporated instead of mZP2^5,12^.

Analysis of native ZP material suggests that the filament cross-linking function of ZP1 is also conserved in human^18,19^; on the other hand, the biological role of ZP4 (previously referred to as ZPB^8,20,21^) remains unknown^22^. However, the two proteins are structurally related by both containing a trefoil domain immediately before their ZP module^23,24^; moreover, ZP1 and ZP4 have been suggested to harbour a single ZP-N-like domain at their N-termini^25,26^. The possible function of this putative element – which is not found in fish homologues of ZP1^27^ – remains nonetheless unclear, considering that multiple copies of isolated ZP-N domains at the N-terminus of ZP2 as well as mollusk vitelline envelope receptor for lysin (VERL) have been shown to regulate sperm binding^28,29^.

Notably, a ZP-N signature can also be recognized at the N-terminus of the avian homologues of ZP1 and ZP4^25,26^. In the case of the former, which together with ZP3 as well as a different peripherally associated subunit (ZPD) is the major constituent of the bird VE^30,31^, the putative N-terminal ZP-N domain is separated from the trefoil domain by a P/Q-rich repeated sequence region that is also conserved in fish and reptilian homologues of the protein^27,30,32^. Moreover, unlike ZP3 and ZPD, which are secreted by the granulosa cells surrounding the egg, avian ZP1 is produced in the liver and reaches the oocyte via the blood circulation^32^. Interestingly, the soluble precursor of ZP1 is monomeric, suggesting that the protein only forms intermolecular disulphides upon incorporation into the growing VE^33^. On the other hand, avian ZP4 is synthesized by the ovary and only expressed in limited amounts during the early stage of folliculogenesis, so that it remains largely localized within the germinal disc region of the eggs where ZP2 is also found^34^.

Because *Zp2*^−/−^ or *Zp3*^−/−^ mice are completely infertile^13–15^ whereas *Zp1*^−/−^ animals are subfertile^16^, only mZP2 and mZP3 are thought to be crucial for mouse fertility; similarly, although a contribution of hZP1 and hZP4 to human sperm binding cannot be completely ruled out^35^, this possibility is not supported by studies of hZP1 “rescue” mice^36^ or transgenic animals expressing hZP4^22^. As a result, the large majority of previous biochemical work focused on ZP2 and ZP3, which have also been investigated by X-ray crystallography^26,29,37,38^. On the contrary, there is no structural information on either ZP1 or ZP4, and the hypothesis that these proteins share a similar function due to their common sequence features remains speculative. Against this background, the recent identification of different ZP1 mutations in infertile patients^39–43^ argues for a much more important role of this glycoprotein in human reproduction than previously recognized. To gain molecular insights into the biological function of ZP1 and its possible relation with that of ZP4, we undertook a biochemical and structural investigation that began with the analysis of a reported case of primary female infertility associated with a frameshift mutation in the human *ZP1* gene (I390fs404X)^39^.

## Results

### Effect of infertility-linked *ZP1* mutation I390fs404X on the secretion of human ZP subunits

Although it was shown that oocytes from homozygous women carrying the *ZP1* I390fs404X mutation lack of a ZP^39^, the effect of the frameshift at the protein level was not investigated. To address this point, we used human embryonic kidney (HEK) 293T cells to compare the expression of the mutant protein, which is truncated shortly after the ZP-N domain (hZP1Mut), to that of wild-type hZP1 (Fig. 1a). This showed that, whereas hZP1 is secreted into the medium as a mixture of monomer and disulphide-bonded dimer, secretion of hZP1Mut is completely abolished (Fig. 1b, left panel). Although the expression of hZP1Mut is lower than that of hZP1, the truncated mutant protein can be detected in the cell lysate (Fig. 1b, right panel). To investigate whether intracellularly-retained hZP1Mut affects secretion of the other human ZP subunits (hZP2-4), we performed co-expression experiments (Fig. 1c, d). As shown in Fig. 1c, secretion of hZP1 increases in the presence of hZP2-4, but co-expression with the latter does not rescue secretion of hZP1Mut. Notably, the secretion levels of hZP2, hZP3 or hZP4 are comparable upon co-expression with either hZP1 or hZP1Mut (Fig. 1d). This data suggests that the human I390fs404X mutation does not affect ZP biosynthesis by interfering with the secretion of the other ZP subunits^39^, but rather by affecting the filament cross-linking function of ZP1.

**Fig. 1.**
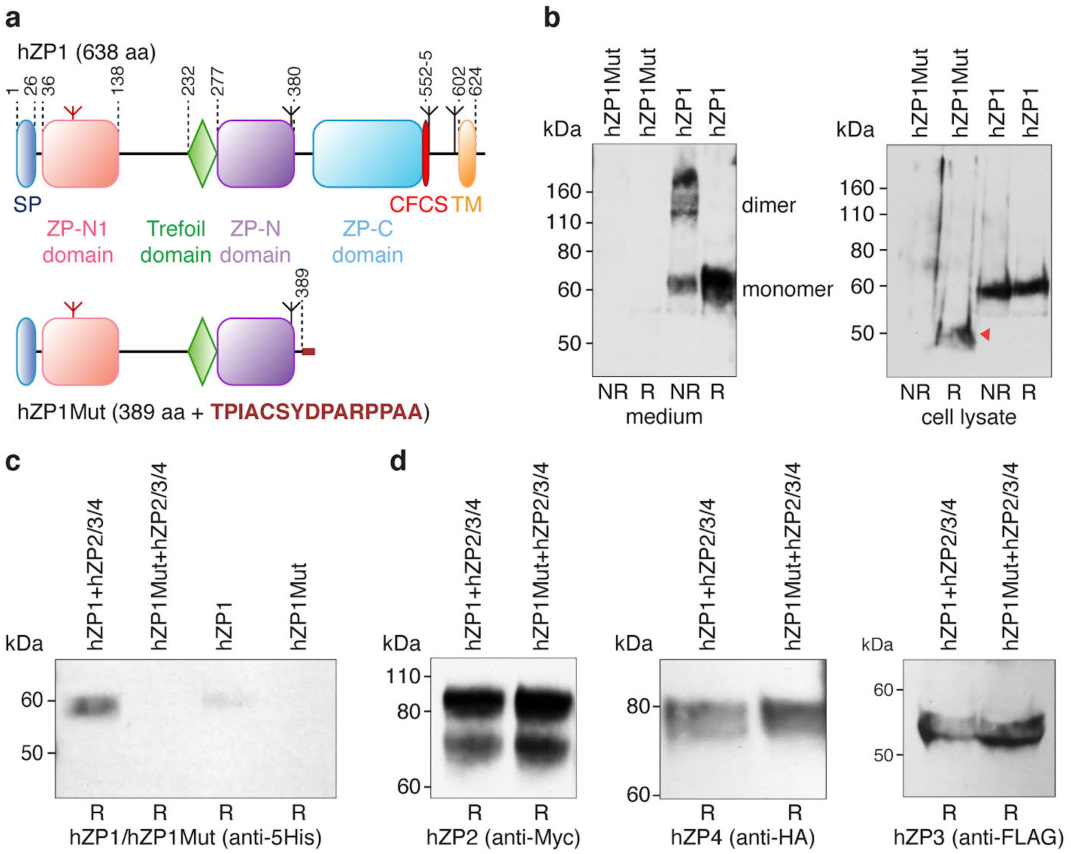
Infertility-associated *ZP1* mutation I390fs404X impairs hZP1 secretion but does not affect co-expressed hZP2, hZP3 and hZP4. **a**, Domain architecture of wild-type and mutant hZP1. SP, signal peptide; CFCS, consensus furin cleavage site; TM, transmembrane domain. Inverted tripods mark N-glycosylation sites, with the conserved N-glycan of the ZP-N1 domain coloured red. Domain boundaries are indicated, and the 15-residue C-terminal extension resulting from the frameshift mutation is highlighted in dark red. **b**, Anti-5His immunoblot of medium (left; 2.4 mL/lane) and cell lysate (right; 0.3 mL/lane) samples from HEK293T cells expressing hZP1 constructs show that hZP1Mut is retained intracellularly (red arrow). NR, non-reducing conditions; R, reducing conditions. **c**, Co-expression with human ZP2-4 increases secretion of wild-type hZP1 but does not rescue secretion of hZP1Mut (240 μL medium/lane). **d**, The secretion levels of hZP2, hZP3 and hZP4 do not change upon co-expression with either hZP1 or hZP1Mut (240 μL medium/lane). Source data are provided as a Source Data file.

### Identification of the ZP1 region mediating ZP filament cross-linking

To determine which part of ZP1 is responsible for introducing cross-links in the ZP, we took advantage of the natural abundance of this subunit in the chicken VE – where it also forms intermolecular disulphides – and the fact that it is specifically degraded upon sperm penetration^44^. Incubation of native VE material with a sperm protease extract releases a soluble ∼21 kDa N-terminal fragment of chicken ZP1 (cZP1) that exists both as a monomer and an intermolecularly disulphide-bonded homodimer (Fig. 2a-f). Accordingly, a recombinant construct corresponding to the predicted ZP-N1 domain of cZP1 (cZP1-N1) is secreted both as a monomeric and a disulphide-bonded dimer, whereas a protein encompassing the trefoil domain and C-terminal ZP module of cZP1 (cZP1_L600-R958_; Fig. 2a) does not form intermolecular disulphides (Fig. 2g). Thus, by recapitulating the oligomeric status of full-length cZP1 (Fig. 2h), cZP1-N1 harbours its cross-linking function. Importantly, this conclusion can be extended to the mammalian homolog of the protein because, mirroring full-length hZP1 (Fig. 1b, left panel), a construct corresponding to hZP1-N1 is also secreted as a homodimer linked by intermolecular disulphide(s) (Fig. 2i). Moreover, size-exclusion chromatography with multi-angle static light scattering (SEC-MALS) analysis of a construct including the trefoil domain and ZP module of hZP1 (hZP1_A195-P590_; Fig. 1a) excludes the possibility that this region of the molecule contributes to its cross-linking function via non-covalent interactions (Fig. 2j).

**Fig. 2.**
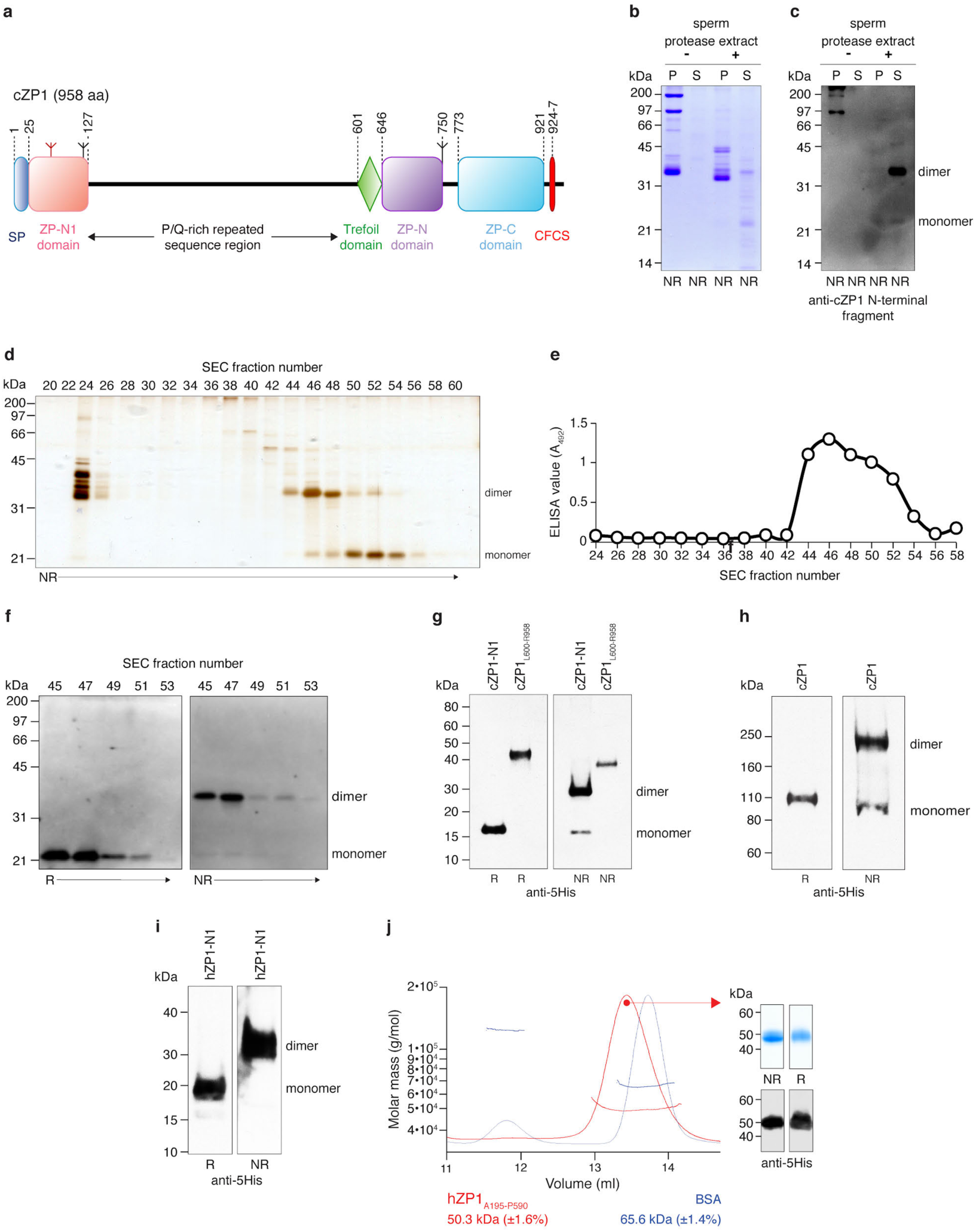
The cross-linking function of ZP1 maps to its N-terminal ZP-N1 domain. **a**, Domain architecture of cZP1, with feature names abbreviated as in Fig. 1a. **b**, Coomassie-stained SDS-PAGE analysis of insoluble (pellet; P) and soluble (S) fractions of native chicken egg coat material that was either untreated or incubated with a sperm protease extract prepared as described previously^44^. **c**, Immunoblot analysis using anti-cZP1 N-terminal fragment polyclonal^57^ of egg coat fractions treated with sperm extract confirms the presence of cZP1-N1 in the soluble fraction. **d**, Silver-stained SDS-PAGE analysis of a size-exclusion chromatography (SEC) separation of the sperm protease-solubilized egg coat material. **e**, Fractions 44-54 contain cZP1-N1 as evidenced by ELISA analysis using anti-cZP1 N-terminal fragment polyclonal^57^. Bands corresponding to the dimer and monomer of cZP1-N1 released from the egg coat were in-gel digested with chymotrypsin and trypsin, followed by MALDI-MS/MS to confirm their protein sequences. **f**, Immunoblot analysis of SEC fractions 45-53 indicates that the ∼35 kDa protein in these fractions is a disulphide-linked dimer of the ∼21 kDa protein containing cZP1-N1. **g-h**, Anti-5His immunoblot of recombinant cZP1-N1 and cZP1_L600-R958_ (g), as well as full-length cZP1 (h). Only cZP1-N1 forms a covalent homodimer like full-length cZP1. **i**, Anti-5His immunoblot analysis shows that secreted hZP1-N1 also forms a covalent homodimer. **j**, SEC-MALS analysis shows that purified hZP1_A195-P590_ produced in HEK293S cells has a molecular mass of ∼50 kDa (red profile), consistent with the calculated mass of a monomeric species carrying 2 GlcNac residues (43 kDa). Calibration was carried out using BSA, whose molecular mass detection is shown in blue. Source data are provided as a Source Data file.

### Three-dimensional structures of the cZP1-N1 cross-link

To establish the molecular basis of ZP1 cross-linking, we investigated the cZP1-N1 homodimer by X-ray crystallography. cZP1-N1 is chemically heterogeneous due to the presence of two N-glycosylation sites (N65 and N121), only the first of which is highly conserved and also found in hZP1 (Supplementary Fig. 1a). Accordingly, whereas mutation of cZP1-N1 N65 or hZP1-N1 N76 drastically reduces protein secretion, a cZP1 N121Q mutant is secreted as well as the wild-type and – because of its higher homogeneity – was used for crystallization trials in parallel with hZP1-N1 (Fig. 3a). Whereas the latter did not crystallize, we could grow crystals of cZP1-N1 expressed in human embryonic kidney (HEK) 293S GnTI-cells and treated with Endoglycosidase (Endo) H (Fig. 3b), as well as the same protein expressed in HEK293T cells (Fig. 3c). The structure of deglycosylated cZP1-N1 was determined by gold- as well as zinc-single-wavelength anomalous dispersion (SAD) phasing. The resulting model was used to solve by molecular replacement the structure of glycosylated cZP1-N1 (Fig. 3d-g and Table 1).

**Fig. 3.**
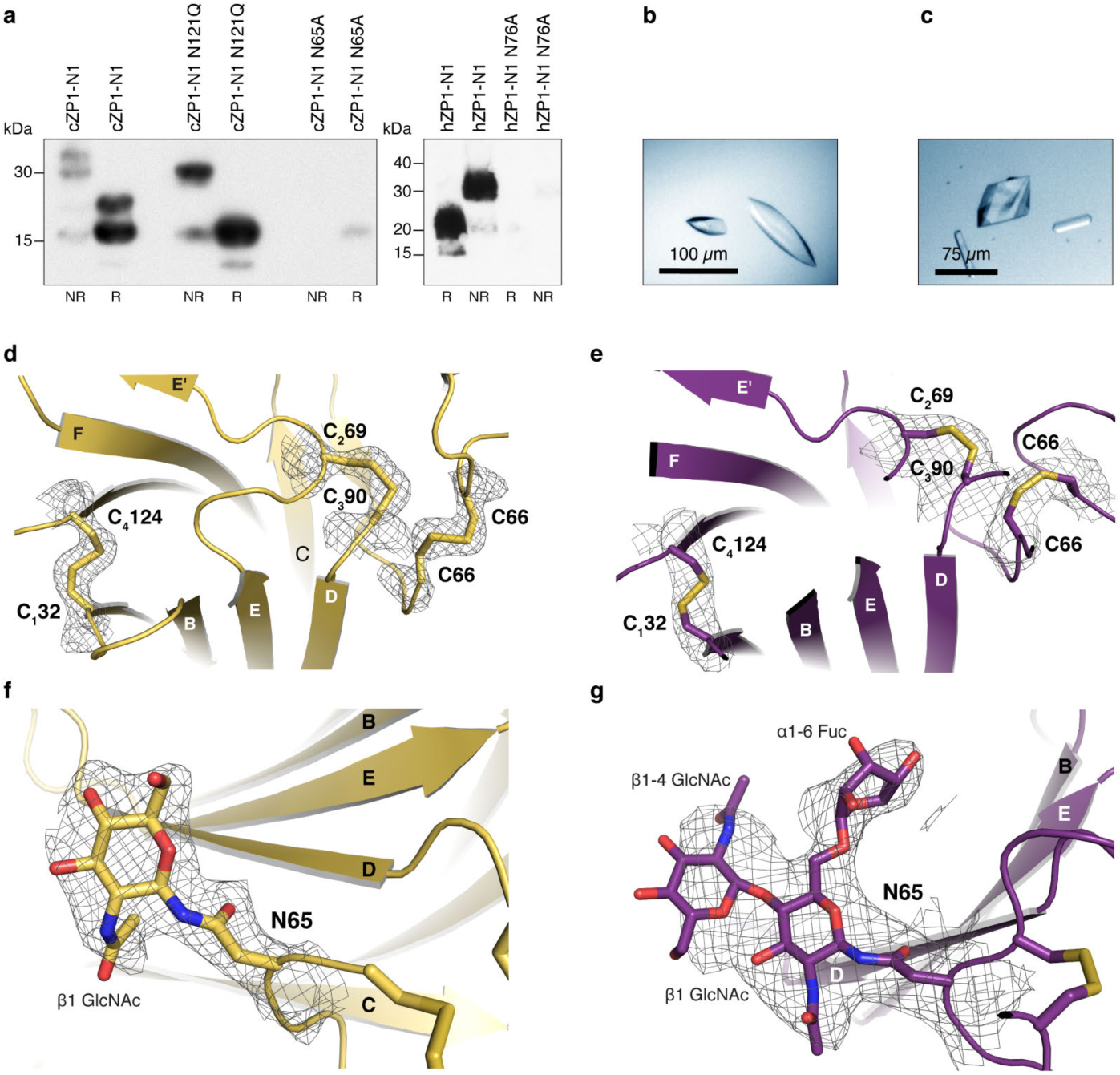
Structure determination of Endo H-deglycosylated and glycosylated cZP1-N1. **a**, Anti-5His immunoblot shows that mutation of N-glycosylation site N65, but not N121, significantly reduces the expression of cZP1-N1 (left panel). Similarly, hZP1-N1 with a N76A mutation is essentially not secreted (right panel). All lanes correspond to 20 μL medium. **b**, Crystals of Endo H-deglycosylated cZP1-N1. **c**, Crystals of glycosylated cZP1-N1. **d-e**, Details of the Endo H-deglycosylated (d) and glycosylated (e) cZP1-N1 structures, depicted in cartoon representation with canonical ZP-N domain disulphides and the C66 cross-link shown as sticks. Relevant portions of the *2mFo-DFc* electron density maps contoured at 1.0 σ are shown as black meshes. **f-g**, Detailed views of the N65-linked glycans of Endo H-treated (f) and glycosylated cZP1-N1 (g). Electron density maps are depicted as in panels d-e. Source data are provided as a Source Data file.

**Table 1.** X-ray data collection and refinement statistics

|  | **cZP1 ZP-N1**<br>**(HEK293S)**<br>**[PDB: 6GF6]** | **cZP1 ZP-N1 Zn<sup>2+</sup>**<br>**(HEK293S)**<br>**[PDB: 6GF7]** | **cZP1 ZP-N1**<br>**(HEK293T)**<br>**[PDB: 6GF8]** |
| --- | --- | --- | --- |
| **Data Collection** |  |  |  |
| Resolution range (Å) | 37.0-2.30 (2.40-2.30) | 53.3-2.70 (2.82-2.70) | 24.7-3.10 (3.24-3.10) |
| Space group | *P* 4<sub>1</sub> 2<sub>1</sub> 2 | *P* 4<sub>1</sub> 2<sub>1</sub> 2 | *C* 2 2 2<sub>1</sub> |
| Unit cell |  |  |  |
| *a*, *b*, *c* (Å) | 75.98, 75.98, 101.88 | 75.41, 75.41, 100.78 | 90.77, 125.73, 105.00 |
| $\alpha$, $\beta$, $\gamma$ (°) | 90, 90, 90 | 90, 90, 90 | 90, 90, 90 |
| Total reflections | 267258 (11835) | 92642 (6475) | 45635 (5509) |
| Unique reflections | 13549 (1420) | 15254 (1896) | 11096 (1326) |
| Multiplicity | 19.7 (8.3) | 6.1 (3.4) | 4.1 (4.2) |
| Completeness (%) | 97.9 (84.5) | 99.9 (99.2) | 98.7 (96.4) |
| Mean *I*/$\sigma$(*I*) | 23.81 (1.49) | 13.63 (1.06) | 10.49 (1.18) |
| Wilson B-factor (Å<sup>2</sup>) | 38.90 | 70.30 | 111.38 |
| R<sub>meas</sub> (%) | 11.6 (149.8) | 10.3 (162.7) | 9.5 (111.2) |
| R<sub>pim</sub> (%) | 2.6 (49.5) | 4.1 (85.2) | 4.4 (51.9) |
| CC<sub>1/2</sub> | 1.00 (0.55) | 1.00 (0.51) | 1.00 (0.64) |
| CC* | 1.00 (0.84) | 1.00 (0.82) | 1.00 (0.89) |
| **Refinement** |  |  |  |
| Reflections | 13547 (1419) | 15254 (1891) | 11096 (1319) |
| Free reflections | 1349 (137) | 1528 (188) | 1070 (122) |
| R<sub>work</sub> (%) | 19.22 (29.53) | 22.18 (36.53) | 21.04 (38.88) |
| R<sub>free</sub> (%) | 22.92 (31.86) | 27.18 (43.14) | 23.52 (43.51) |
| CC<sub>work</sub> | 0.960 (0.746) | 0.927 (0.610) | 0.947 (0.816) |
| CC<sub>free</sub> | 0.935 (0.743) | 0.905 (0.438) | 0.930 (0.696) |
| Number of non-H atoms | 1834 | 1723 | 1736 |
| macromolecules | 1712 | 1660 | 1638 |
| ligands | 34 | 37 | 98 |
| solvent | 88 | 26 | 0 |
| Protein residues | 216 | 208 | 217 |
| RMS deviations |  |  |  |
| bonds (Å) | 0.005 | 0.004 | 0.003 |
| angles (°) | 0.72 | 0.72 | 0.65 |
| Average B-factor | 47.54 | 74.13 | 140.69 |
| macromolecules | 47.29 | 73.51 | 138.49 |
| ligands | 68.86 | 97.41 | 177.50 |
| solvent | 44.32 | 80.40 | - |

Despite very low sequence identity, the protein adopts the same modified immunoglobulin (Ig)-like fold of the ZP2-N1 and ZP3 ZP-N domains^5,26,29,37^, including two intramolecular disulphides with 1-4, 2-3 connectivity (C_1_32-C_4_124 and C_2_69-C_3_90), a Tyr residue located in β-strand F next to C_4_ and an E’ strand (Fig. 4). However, the cd loop of cZP1-N1 includes an additional β-strand (C’) that forms a β-hairpin with strand C. Together with invariant V63, which interacts hydrophobically with the CD face, this positions an additional Cys (C66) so that it can form an intermolecular disulphide with the same residue from another molecule (Figs. 4 and 5).

**Fig. 4.**
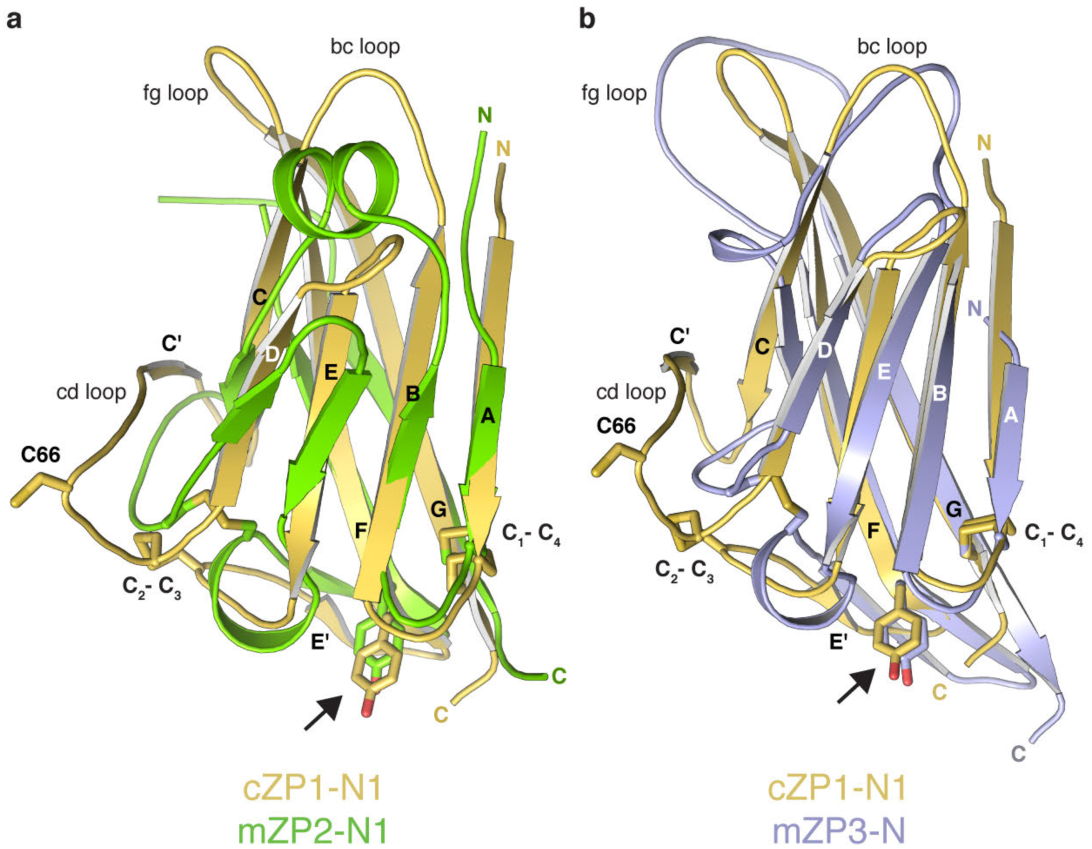
Structural comparison of ZP-N domains from vertebrate ZP1, ZP2 and ZP3. **a**, Superimposition of cZP1-N1 (chain B of the highest resolution structure, PDB 6GF6) and mZP2-N1 (PDB 5II6), with a root-mean-square deviation (RMSD) of 2.1 Å over 74 aligned residues (13% sequence identity). Cross-linking C66, conserved disulphides and β-strand F Tyr (arrow) are shown as sticks. **b**, Superimposition of cZP1-N1 and mZP3-N (chain A of PDB 5OSQ), with a RMSD of 2.1 Å over 75 aligned residues (11% sequence identity). Note how the bc and fg loops of cZP1-N1 are significantly shorter than those of mZP3-N.

**Fig. 5.**
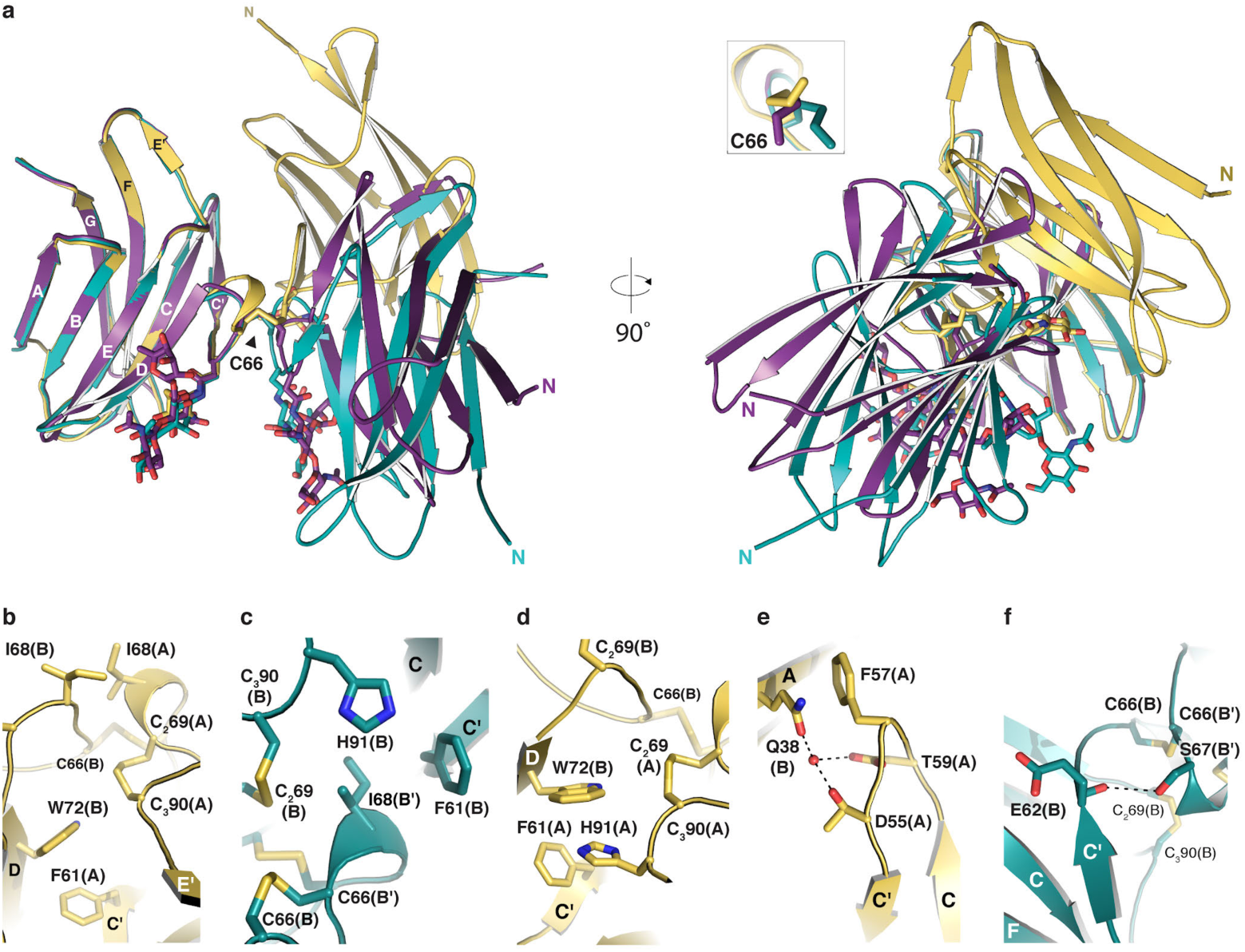
Conformational variability of the ZP1 cross-link. **a**, Different relative arrangements of the subunits of the Endo H-treated (chains A/B, gold) and glycosylated (chains A/A’, violet purple; chains B/B’, teal) cZP1-N1 homodimers. The N-glycan attached to conserved N65 and the C66 cross-link are shown as sticks. **b-f**, Details of the asymmetric and symmetric interfaces of the Endo H-treated and glycosylated homodimers, respectively. Structures are coloured as in panel a, with selected amino acids shown as sticks and hydrogen bonds indicated by dashed lines.

Remarkably, although they all contain the same C66-C66 cross-link, the moieties of the cZP1-N1 homodimers observed in our crystal forms adopt three distinct relative orientations (Fig. 5a). These correspond to an asymmetric interface in the case of deglycosylated cZP1-N1 (whose crystallographic asymmetric unit contains a homodimer) and two related but not identical interfaces in the crystal of glycosylated cZP1-N1 (whose asymmetric unit also contains two copies of the protein (chains A and B) that, however, make intermolecular cross-links with symmetry-related copies of themselves (A’ and B’)). All the cZP1 cross-links have small interface areas that range from 330/412 Å^2^ (A-A’/B-B’ dimers of glycosylated cZP1-N1) to 575 Å^2^ (deglycosylated cZP1-N1 dimer). Moreover, in addition to C66, they involve a common number of invariant or highly conserved residues including F61, N65, S67, the C_2_69-C_3_90 intramolecular disulphide and H91, as well as I68 (conserved in cZP1, hZP1 and mZP1) (Supplementary Fig. 1a). Distinct sets of interactions mediated by these residues underlie the different homodimer interfaces: for example, in the deglycosylated cZP1-N1 homodimer, C’ strand F61(A) interacts aromatically with W72(B) and I68(A) packs against I68(B) (Fig. 5b); on the other hand, in both glycosylated cZP1-N1 homodimers, the latter interaction is missing because I68 interacts hydrophobically with F61 of the opposite subunit (Fig. 5c). Similarly, while H91(A) stacks against W72(B) in the deglycosylated cross-link structure (Fig. 5d), H91 faces I68 in both of the glycosylated cross-links (Fig. 5c).

Additional contacts between residues in the region that encompasses β-strands A-B and residues in the cc’ loop stabilizes the asymmetric structure of the deglycosylated cZP1-N1 cross-link; in particular, invariant Q38(B) stacks against highly conserved F57(A) and also makes water-mediated hydrogen bonds with the side chains of T59(A) and invariant D55(A) (Fig. 5e). In the case of glycosylated cZP1-N1, on the other hand, the relatively more extensive B/B’ molecule interface is stabilized by a long-range hydrogen bond between the side-chain oxygen of conserved S67 and the main-chain carbonyl oxygen of E62 (Fig. 5f).

### The intermolecular disulphide is essential for ZP1-N1 homodimerization in chicken, human and mouse

Consistent with the evolutionary conservation of C66 (Fig. 6a and Supplementary Fig. 1a) and the observation that the homodimer interfaces observed in the cZP1-N1 crystals bury a limited fraction of surface area, SDS-PAGE analysis of SEC fractions show that mutation of C66 abolishes protein homodimerization (Fig. 6b, c). Similarly, mutation of the corresponding residues of hZP1-N1 and mZP1-N1 (C77 and C69, respectively) completely disrupts their ability to homodimerize (Figs. 6d, e and 6f, g). Notably, whereas the conditioned medium of cells expressing cZP1-N1 contains a small fraction of monomeric protein in addition to the prevalent homodimeric species, secreted hZP1-N1 and mZP1-N1 are entirely dimeric (compare Fig. 6c with Fig. 6e, g). Together, these data indicate that formation of the conserved intermolecular disulphide is required for ZP1-N1 homodimerization in both birds and mammals, although the efficiency of this process may vary between the two.

**Fig. 6.**
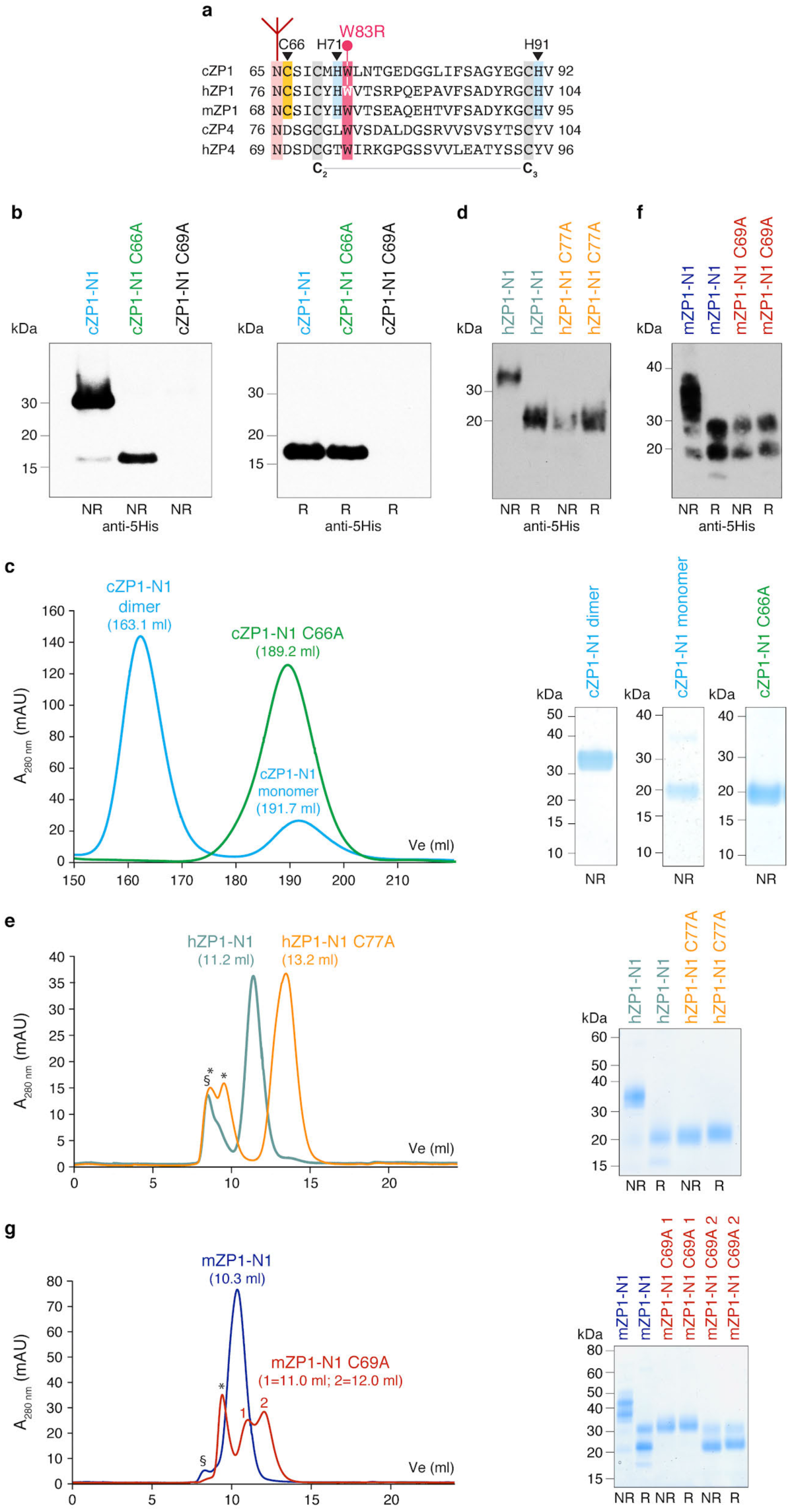
The conserved non-canonical Cys of ZP1-N1 mediates protein homodimerization in avian and mammalian species. **a**, Sequence alignment of selected ZP1/4 homologues. The conserved N-glycosylation site and non-canonical Cys of ZP1 are highlighted in pink and yellow, respectively, whereas canonical ZP-N domain Cys 2 and 3 are shaded grey. hZP1 W83, mutated in infertile patients (Fig. 7), and conserved ZP1-N1 His residues involved in Zn^2+^ binding (Fig. 8) are shaded in red and cyan, respectively. **b**, Immunoblots with anti-5His monoclonal show that, whereas mutation of C_2_69 abolishes protein expression, a C66A mutation does not affect secretion of cZP1-N1 but completely prevents its covalent homodimerization. **c**, Coomassie-stained SDS-PAGE analysis of SEC experiments performed with a HiLoad 26/600 Superdex 75 pg column confirms that immobilized metal affinity chromatography (IMAC)-purified wild type cZP1-N1 consists of two species, corresponding to dimeric and monomeric protein. On the contrary, cZP1-N1 C66A is entirely monomeric. **d-g**, Parallel analyses show that homodimerization of the human (d, e) and mouse (f, g) ZP1-N1 domains is mediated by C77 and C69, respectively. SEC runs were performed using a Superdex 75 Increase 10/300 GL column; void volume and contaminant peaks are indicated by § and *. As supported by deglycosylation experiments with PNGase F (Supplementary Fig. 2), heterogeneous glycosylation of the two sequons of mZP1-N1 (N49 and N68)^17^ results in two distinct bands on SDS-PAGE. Source data are provided as a Source Data file.

### Effect of fucosylation and infertility-associated hZP1 mutation W83R on ZP1-N1 crosslinking

C66 is part of an almost invariant sequon (Fig. 6a and Supplementary Fig. 1a) whose glycosylation is crucial for cZP1-N1 and hZP1-N1 secretion (Fig. 3a). Interestingly, heterogeneous core fucosylation of native mouse and human ZP carbohydrates was observed by mass spectrometry^45,46^, which also suggests that the same heterogeneity exists in native cZP1-N1 (Supplementary Fig. 3); however, the relative domain arrangement found in the crystals of deglycosylated cZP1-N1 (Fig. 5a) would not be compatible with the presence of an α1-6-linked fucose (Supplementary Fig. 4). These observations suggest that the structure of the conserved N-glycan of ZP1 could modulate the conformation of ZP filament cross-links by favouring a specific type of ZP1/ZP1 interface.

In the structure of glycosylated cZP1-N1 as well as chain A of the deglycosylated protein, the cd loop containing C66 packs against W72, a highly conserved residue in β-strand D (Fig. 6a and Supplementary Fig. 1a). In chain A of glycosylated cZP1-N1, this residue also stacks against the ring of the α1-6-linked fucose (Fig. 7a); on the contrary, as mentioned above, W72(B) stacks against H91(A) in the asymmetric interface of the deglycosylated homodimer (Fig. 5d). Notably, a mutation of the corresponding amino acid of human ZP1 (W83R) was recently found in an infertile patient lacking the ZP (H. Zhao, personal communication and reference 40). To investigate the effect of this pathogenic substitution, we compared the expression of a hZP1-N1 W83R mutant to that of the wild-type protein. Consistent with an important role of W72 in the folding and dimerization of cZP1-N1, this experiment showed that W83R not only decreases the secretion of hZP1-N1, but also significantly reduces its ability to form disulphide-bonded homodimers (Fig. 7b). This suggests that the mutation causes lack of the ZP by hindering human ZP filament cross-linking.

**Fig. 7.**
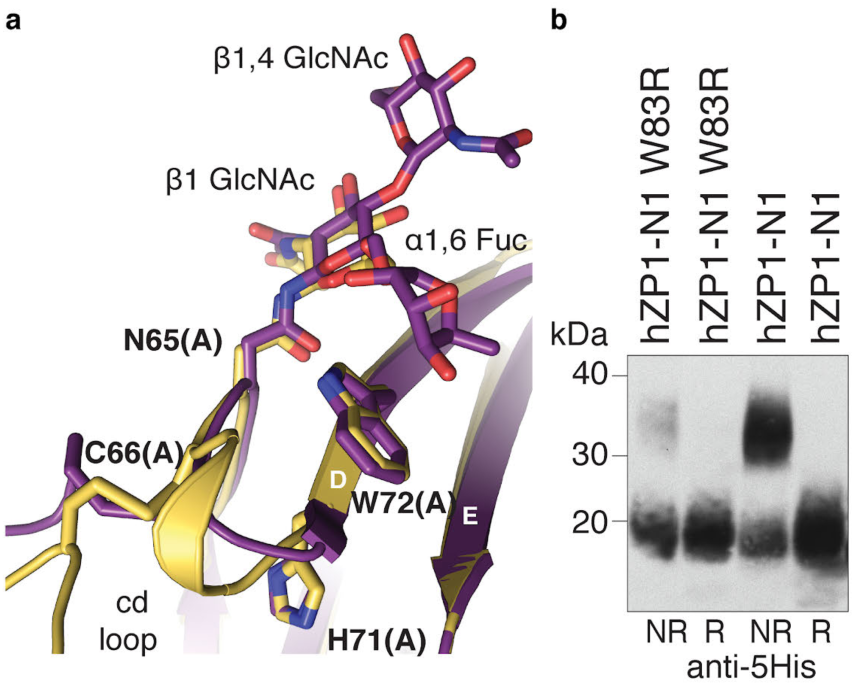
Infertile patient hZP1 mutation W83R affects secretion and homodimerization of hZP1-N1. **a**, W72 stacks between the cd loop and the N65 glycan α1,6 fucose. **b**, Anti-5His immunoblot of hZP1-N1 W83R mutant (20 μL medium) compared to wild-type (10 μL medium) 72 h post-transfection. Source data are provided as a Source Data file.

### Two conserved Zn^2+^-binding His of cZP1-N1 are important for hZP1 cross-link formation

Examination of the zinc-SAD data used for phasing revealed that asymmetric interaction of the cZP1-N1 homodimer moieties creates a negatively charged pocket where H71(A) coordinates a Zn^2+^ ion that faces the C_2_69(A)-C_3_90(A) intramolecular disulphide, as well as the C66 cross-link (Fig. 8a). The anomalous difference map of the data also shows a peak near H71(B) but, consistently with the much lower accessibility of the latter due to crystal packing, this site is weaker than the previous (∼6 σ vs 11 σ). In the native electron density maps of cZP1-N1, the Zn^2+^ is missing but the same His residue, together with closely located H91, is part of the homodimer interface in both its asymmetric and symmetric conformations. In the former case, the pocket is occupied by a molecule of glycerol from the cryoprotectant solution and H91(A) forms an aromatic residue cluster with W72(B) and F61(A) (Fig. 5d and Fig. 8b). The imidazole ring of H71(A) stacks against the C_2_69(A)-C_3_90(A) disulphide and is also hydrogen-bonded to main chain carbonyl group of C_3_90(A); at the same time, the main chain atoms of H71(A) stabilize the intermolecular cross-link by hydrogen bonding to the backbone of C66(A) (Fig. 8b). In the case of the symmetric interface, H71 is also H-bonded to C_3_90 and C66 and, together with H91 from the same moiety of the homodimer, lines the wall of a cavity whose edge interacts hydrophobically with I68 from the other subunit (Fig 8c). Notably, the two His residues and the W72 are almost completely conserved in ZP1 homologs from other species, including mouse and human (Fig. 6a and Supplementary Fig. 1a).

**Fig. 8.**
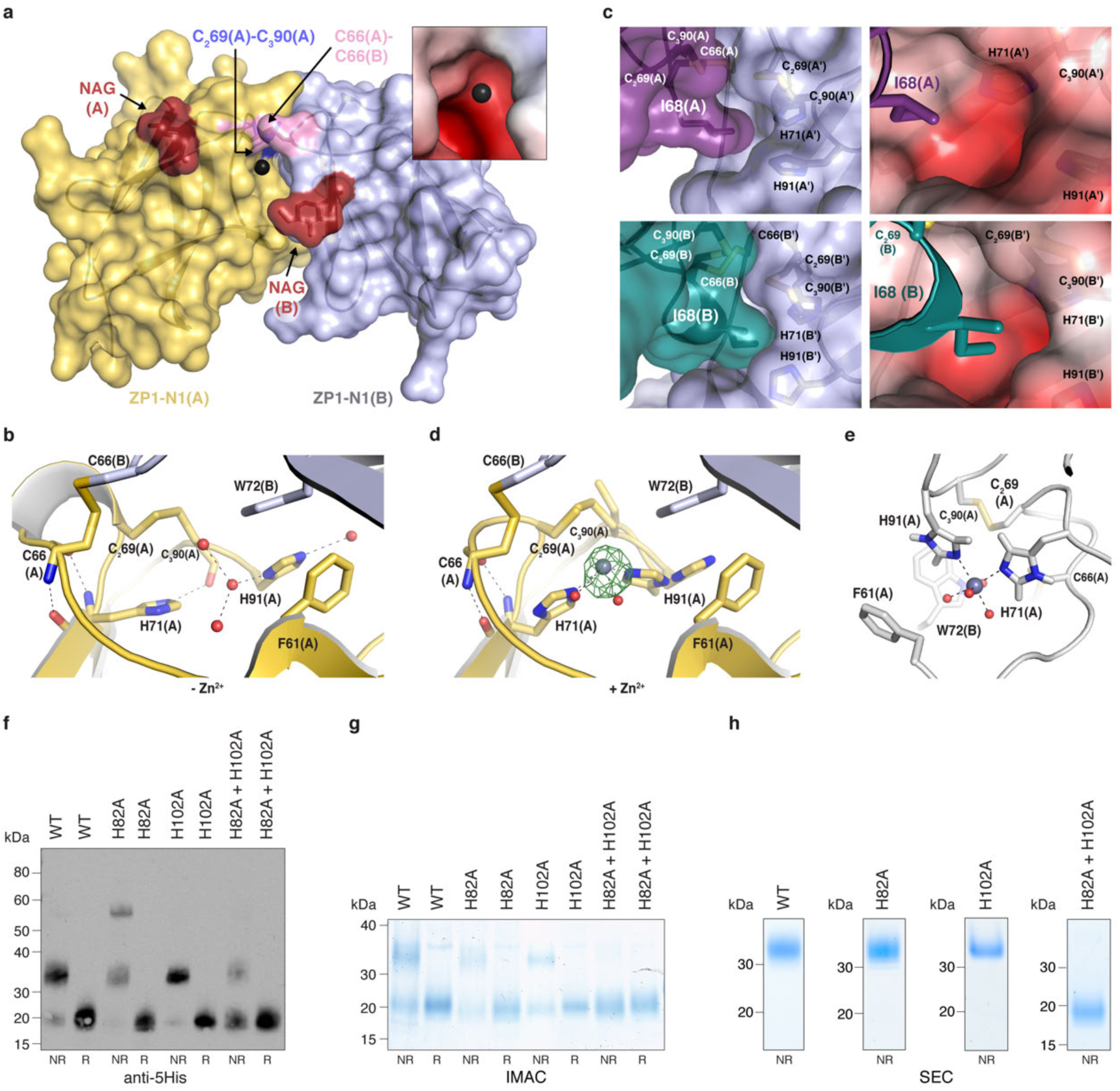
hZP1 cross-link formation depends on a conserved His pair that can bind Zn^2+^ in cZP1-N1. **a**, Surface representation of the structure of the deglycosylated cZP1-N1 homodimer from a crystal soaked with Zn(OAc)_2_, highlighting the position of the Zn^2+^ ion bound to H71(A) and H91(A) (black sphere) relative to the protein subunits, two proximal disulphide bonds and N-acetylglucosamine (NAG) residues attached to N65. The A subunit of the homodimer is coloured yellow according to Fig. 5, whereas subunit B is shown in lilac. The inset shows a detailed view of the region around the Zn^2+^, coloured by electrostatic potential using a gradient from red (−4 kT/e) to blue (+4 kT/e) through white (0 kT/e). **b**, Surface representation of the symmetric interface of glycosylated cZP1-N1 (chain A, top left panel; chain B, bottom left panel), showing the position of H71 and H91 relative to I68. Notably, the H71/H91 pocket of the A’ subunit is more accessible than that of B’, due to apparent flexibility of the relatively small symmetric interface. The right panels, which are rotated by 30 degrees over the Y axis compared to the left ones, show zoomed views of the symmetric interfaces around I68(A) and I68(B), with the surface of the A’ and B’ molecules coloured by electrostatic potential as in panel a. **c-d**, Details of the asymmetric interface of deglycosylated cZP1-N1 as observed in native (c) or Zn^2+^-soaked (d) crystals. Selected amino acids are shown as sticks, with hydrogen bonds indicated by dashed lines. The green mesh in panel d is an anomalous difference map calculated at λ=1.2825 Å and contoured at 6 σ. **e**, 100-ns MD simulation snapshot of the Zn^2+^ binding site shown in panel a. H71 and H91, together with four water molecules, coordinate the zinc ion during the whole simulation time (mean Zn-His N distance 2.22±0.06 Å; mean Zn-H_2_O O distance 2.10±0.05 Å). **f**, Anti-5His immunoblot analysis of wild-type (WT) and His mutant hZP1-N1 constructs secreted by HEK293T cells (100 μL medium). **g-h**, Coomassie-stained SDS-PAGE analysis of IMAC (g) and SEC (h) fractions from the purification of the same constructs as in panel f. The SEC profiles corresponding to panel h are shown in Supplementary Fig. 5. Source data are provided as a Source Data file.

In the Zn^2+^-soaked crystal, where H71(A) adopts a different orientation and is too far from C_3_90 (A) to make an hydrogen bond, the C_2_69(A)-C_3_90(A) disulphide is partially reduced (most likely due to radiation damage) and H91(A) adopts two alternative conformations, one of which no longer stacks against W72(B) and chelates the Zn^2+^ together with H71(A) itself (Fig. 8d). Accordingly, molecular dynamics (MD) simulations suggest that, in solution, the two His residues can stably coordinate a zinc ion together with 4 water molecules (Fig. 8e).

Considering that physiological release of zinc from activated oocytes has been suggested to remodel the mammalian ZP^47,48^, we evaluated the effect of mutating the conserved His residues on the formation of the intermolecular cross-link of hZP1-N1. Whereas mutation of H102 (corresponding to cZP1 H91) does not hinder protein homodimerization, a construct carrying a mutation of H82 (corresponding to cZP1 H71) is secreted as a mixture of disulphide-bonded dimers and tetrameric aggregates; combination of the two mutations, on the other hand, drastically interferes with cross-linking of hZP1-N1 (Fig. 8f-h).

Taken together, these observations indicate that homodimerization of hZP1 depends on the same His residues that form a Zn^2+^-binding site in cZP1-N1.

### Oligomerization properties of human and chicken ZP4-N1

Considering that ZP subunit ZP4 has the same domain architecture as ZP1, its ZP-N1 domain could in principle also harbour a cross-linking function. However, SEC analysis of purified hZP4-N1 (Fig. 9a) as well as pull-down experiments of Myc-tagged hZP4-N1 using an equivalent histidine-tagged construct (Fig. 9b) indicate that the protein is entirely monomeric. On the other hand, parallel experiments using cZP4-N1 indicate that, in the chicken, ZP4 forms a non-covalent homodimer (Fig. 9a, c).

**Fig. 9.**
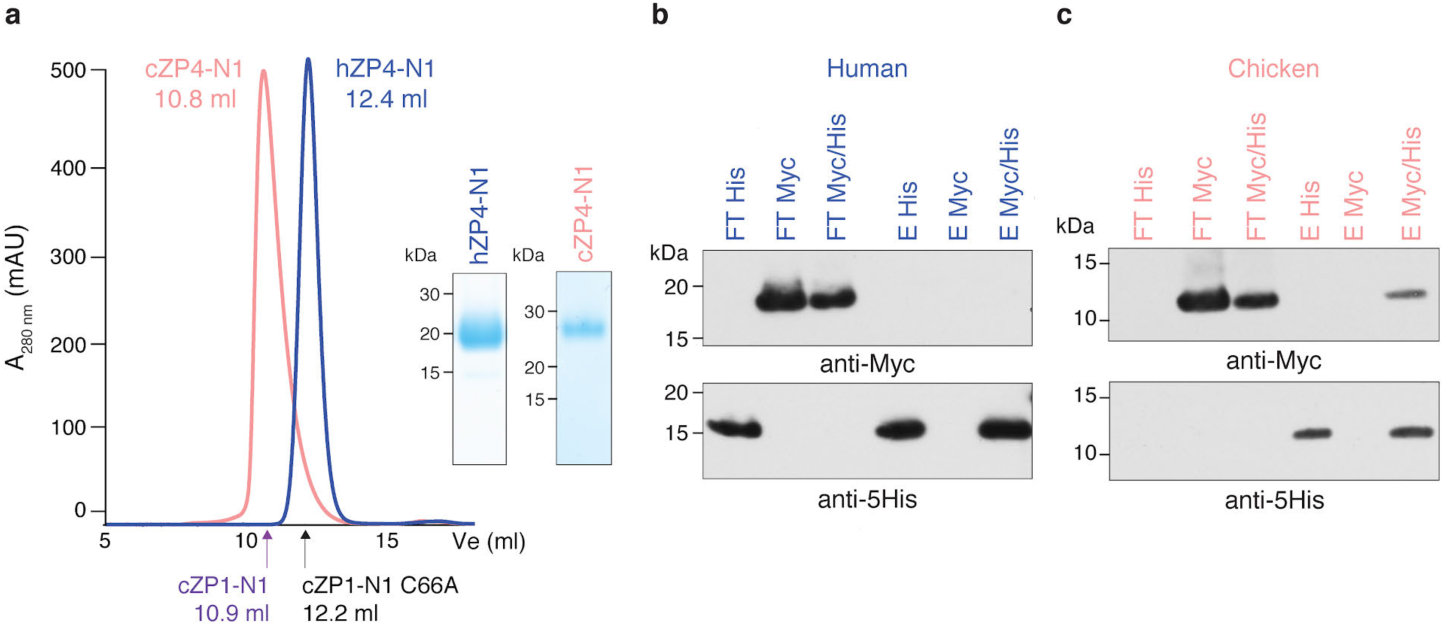
ZP4-N1 forms a non-covalent homodimer in chicken but not human. **a**, SEC analysis of purified hZP4-N1 and cZP4-N1 and non-reducing Coomassie-stained SDS-PAGE analysis of the corresponding peak fractions. Peaks are normalized to 500 mAU, and elution volumes of the cZP1-N1 homodimer and the cZP1-N1 C66A monomer are indicated. **b-c**, Immunoblot analysis in reducing conditions of His-pull-down experiments of co-expressed hZP4-N1-His/Myc or cZP4-N1-His/Myc. Samples were PNGase F-treated before SDS-PAGE. FT, flow-through; E, elution. Source data are provided as a Source Data file.

## Discussion

Although ZP1 remains by far the less studied component of the mammalian ZP, our characterization of the hZP1 I390fs404X mutant protein (Fig. 1) suggests that infertile patients carrying the corresponding mutation lack the ZP due to an impairment of hZP1 secretion that in turn causes absence of filament cross-linking, rather than interference with hZP3 secretion as previously hypothesized^39^. Indeed, even if mutated hZP1 could cause intracellular retention of hZP3 or other ZP subunits, this would only affect a small fraction of the latter. This is because hZP2-4 are expressed in significantly higher amounts than hZP1^21,49,50^ and – in agreement with observations in the mouse^51^ – can be secreted by mammalian cells independently from each other (reference 19, where hZP4 was misannotated as hZP1^8,20,21^). Such a situation would thus be unlikely to cause complete absence of the ZP, also considering that *Zp3*^+/−^ heterozygous mice that lack approximately 50% of ZP3 are still able to assemble a ZP of reduced thickness and, like *Zp2*^+/−^ animals, retain normal fertility^13–15,52^.

Consistent with an essential role of hZP1 and/or hZP4 in human ZP assembly, transgenic mice lacking mouse ZP subunits but expressing only hZP2 and hZP3 cannot form a ZP^22^; moreover, despite its structural similarity to hZP1, hZP4 is clearly not sufficient to rescue ZP biogenesis in hZP1 I390fs404X patients^39^. The latter observation, together with the fact that the trefoil/ZP module region of hZP1 does not form non-covalent homodimers (Fig. 2j), suggest that hZP1-N1 cross-link formation is essential for the assembly of the human ZP and, hence, fertility. Notably, two other *ZP1* variants have been found together with I390fs404X in infertile patients with empty follicle syndrome^41,43^. Similarly to I390fs404X itself, both of these mutations are expected to impair the cross-linking ability of hZP1 by either replacing an exposed residue of hZP1-N1 β-strand C with an additional Cys^41^ or truncating the protein at the level of the hZP1-N1 bc loop^43^. Also taking into account the identification of additional hZP1-truncating variants associated with lack of a ZP^42^, a picture is starting to emerge whereby different constellations of homozygous or compound heterozygous *ZP1* mutations cause primary infertility by hindering human ZP cross-linking and/or oocyte growth. Moreover, in relation with the first aspect, it is apparent that *ZP1* mutations can have a damaging effect not only by completely abolishing ZP cross-linking due to impairment of protein secretion (I390fs404X; Fig. 1), but also by specifically affecting hZP1 residues (such as W83R; Fig. 7) whose structural importance affects cross-link formation. Because of the significant sequence identity between cZP1-N1 and its mouse and human counterparts (53% and 51%, respectively; Supplementary Fig. 1), the crystallographic information reported in this manuscript provides a solid framework for understanding the effect of the aforementioned mutations on human ZP cross-linking, as well as interpreting additional infertility-associated hZP1-N1 variants that will be identified in the future.

From a biological point of view, our structural studies validate the hypothesis^25,26^ that – as also recently seen in the case of ZP2^29^ – the N-terminal region of ZP1 (and, by extension, ZP4) also adopts a ZP-N fold (Fig. 4). However, the cZP1-N1 domain also contains a short C’ β-strand (R60-E62) that contributes to its homodimerization (Fig. 5b-d and f). Because such a strand had previously only been found in ZP-C domains as part of a C’-C” β-hairpin^5^, its presence within cZP1-N1 brings additional support to the idea that ZP-N and ZP-C originated by duplication of an ancestral Ig-like domain^37^.

Most importantly, analysis of the cZP1-N1 cross-link crystals reveals the presence of different homodimer interfaces, stabilized by alternative interactions involving a set of largely conserved residues (Fig. 5 and Supplementary Fig. 1). This suggests that the invariant intermolecular disulphide of ZP1 – whose mutation abolishes the homodimerization of chicken, mouse and human ZP1-N1 (Fig. 6) – allows the protein to introduce a cross-link that is not only mechanically and chemically resistant, but also able to join ZP filaments with highly variable relative orientations. By requiring multiple interactions to be comparably strong, a non-covalent interface would be much less plastic and thus enforce significant constraints on the geometry of filament cross-linking. Interestingly, sequence covariation considerations support the interfaces observed in the cZP1-N1 crystals by showing that, in some species from the suborder Serpentes, both members of the H91(A)/W72(B) stacking pair (and in one case also F61(A)) are replaced by non-aromatic residues, whereas I68 is substituted with a more hydrophobic Phe; at the same time, the conserved sequon preceding the cross-linking Cys has been replaced by another within the de loop (Supplementary Fig. 1b). As a result of this kind of changes, one type of homodimer interface may be favoured over the other in particular subsets of species.

Notwithstanding the molecular contortionism of the cZP-N1 homodimer, structural considerations suggest that core fucosylation of the essential N-glycan that immediately follows the intermolecular disulphide (Fig. 3 and Supplementary Fig. 1) may favour the symmetric interface of the cZP1-N1 cross-link over its more extensive asymmetric alternative (Supplementary Fig. 4). Because cortical granule exocytosis releases small amounts of α-fucosidase^53^, an enzyme that has also been detected in the mammalian oviductal and uterine fluid^54,55^, defucosylation of hZP1 could contribute to post-fertilization remodelling of the ZP by altering the mechanical properties of its cross-links.

The observation that a Zn^2+^ ion can be bound by two His residues of cZP1-N1 (Fig. 8), one of which is invariant (cZP1 H71/hZP1 H82) and the other almost completely conserved (cZP1 H91/hZP1 H102) (Supplementary Fig. 1a), also raises a possible connection with the “zinc spark” response of mammalian oocytes to activation. By co-ordinately releasing intracellular zinc into the extracellular space, this exocytotic event – which has also been described in human^47^ – allows resumption of the cell cycle^56^. At the same time, zinc sparks have been suggested to contribute to the block to polyspermy by inducing physiochemical changes into ZP^48^, but the molecular basis of this effect is unknown. The presence of a conserved Zn^2+^-binding site in ZP1 (Fig. 8d, e), together with the finding that mutation of the two Zn^2+^-binding His severely impairs homodimerization of hZP1-N1 (Fig. 8f-h), suggests that zinc ions released by the zygote could modulate the architecture of the ZP by altering the conformation of the ZP1 cross-links. In agreement with this possibility, which does not exclude the presence of additional binding sites in other ZP subunits^48^, H71 directly stabilizes the cZP1-N1 intermolecular disulphide via two main-chain H-bonds in both asymmetric and symmetric homodimer interfaces (Fig. 8b). H91, on the other hand, stabilizes the asymmetric interface by stacking against W72 from the opposite molecule (Fig. 5d and 8b) whereas, in the case of the symmetric homodimer, contributes to the I68-binding pocket together with H71 (Fig. 5c and left panels of Fig. 8c). Notably, like the Zn^2+^-binding pocket of the asymmetric homodimer (Fig. 8a, inset), the I68-binding pocket also has a negatively charged potential (Fig. 8c, right panels). This suggests that exocytosed zinc may not only affect the asymmetric interface by interfering with the stacking between H91(A) and W72(B) (Fig. 8d), but also alter the symmetric interface by competing with I68. Of direct relevance to these considerations and consistent with the fact that zinc spark-induced ZP architecture changes are triggered by relatively high concentrations of the ion^48^, we do not observe bound Zn^2+^ in native crystals of cZP1-N1 (Fig. 8b).

The finding that hZP4-N1 is entirely monomeric (Fig. 9a, b) explains why – as mentioned above – hZP4 does not rescue ZP formation in infertile patients carrying ZP1 mutations^39^. While mammalian ZP4 could be involved in increasing the thickness of the ZP (reference 28 and Supplementary Table 1), cZP4-N1 is markedly different from its human counterpart in that it is entirely secreted as a homodimer (Fig. 9a, c). This observation supports the recent suggestion that cZP4 functionally substitutes cZP1 in the VE of the immature oocyte, which is assembled before cZP1 starts being expressed^30,34^. Consistent with the role of cZP1 as a major component of the VE of mature oocytes^30,34^, the P/Q-rich repeated region of this subunit (absent in cZP4) may act as a molecular spring that stretches while the egg coat dramatically expands over the course of oocyte maturation. This would allow the cZP1-N1 cross-links to be maintained, until they are released (Supplementary Fig. 3) when penetrating sperm degrades the P/Q-rich region of cZP1^30,44^. While the very low abundance of mammalian ZP1 makes it difficult to determine whether this is also proteolyzed upon sperm penetration, it is interesting to note that ZP1-N1 domains are not found in fish, where sperm penetrates the egg coat through the micropyle. However, fish VE subunits also contain N-terminal P/Q-rich regions that are covalently cross-linked by transglutaminase during post-fertilization hardening^27^.

In summary, our studies provide the first structural information on ZP1 and reveal the molecular basis of egg coat cross-linking from birds to mammals. Considering that the intermolecular cross-link of hZP1 is only formed upon its secretion into the extracellular space (Fig. 1b), the identification of this previously neglected subunit as critical for human fertility also highlights it as a novel possible target for non-hormonal contraception.

## Methods

### DNA constructs

cDNA fragments encoding 6His-tagged hZP1 and hZP1Mut (M1-S389 followed by the sequence TPIACSYDPARPPAA), Myc-tagged hZP2, FLAG-tagged hZP3 and HA-tagged hZP4 were synthesized (GenScript; GeneArt/Thermo Fisher Scientific) and subcloned into pHLsec3^29^. A synthetic construct encompassing cZP1-N1 (cZP1_L25-G149S_) was subcloned into pJ609 (DNA2.0/ATUM) in frame with sequences coding for an IgK signal peptide and a 6His-tag inserted within the C-terminal linker fragment following conserved C_4_124; the same DNA was combined with a PCR fragment amplified from cZP1full/pGEM^57^ to generate a 6His-tagged full-length cZP1 ORF in pSI (Promega). The pHLsec3 construct expressing C-terminally 6His-tagged mZP1-N1_M1-A141_ was generated by PCR, using as template Addgene plasmid 14644 (Mouse ZP1 (JD#264)^58^). Constructs expressing C-terminally His- or Myc-tagged versions of hZP1-N1 (hZP1_M1-A149_), hZP4-N1 (hZP4_M1-T140_) and cZP4-N1 (cZP4_M1-S148_) were generated by PCR from the above constructs or pTargeT/cZP4^34^. cDNAs encoding conserved His mutants of cZP1-N1 and hZP1-N1 were synthesized (ATUM) and cloned into pJ609 and pHLsec3, respectively, as described above; the pHLsec3 construct expressing C-terminally 8His-tagged hZP1_A195-P590_ was generated by overlap extension PCR. Oligonucleotides were ordered from Sigma-Aldrich and constructs were verified by DNA sequencing (Eurofins Genomics).

### Mammalian protein expression

Proteins were transiently expressed as previously described^59^ using HEK293 cultured in serum-free media, with the exception of hZP1_A195-P590_ whose production was carried out using media supplemented with 2% foetal bovine serum (Biological Industries). Co-expression experiments were performed using equivalent ratios of plasmid DNAs; for control single-expression experiments, expression plasmids were supplemented with empty vector DNA (pSI; Promega) to maintain the same ZP subunit/total DNA ratio.

### Protein analysis

For immunoblotting, proteins separated by SDS-PAGE were transferred to a nitrocellulose membrane (GE Healthcare Life Sciences) and probed with primary antibodies anti-5His monoclonal (1:1,000; QIAGEN), anti-Myc monoclonal (1:1,000; Sigma-Aldrich clone 9E10), anti-FLAG monoclonal (1:1,000; Sigma-Aldrich clone M2), anti-HA monoclonal (1:1,000; Sigma-Aldrich clone HA-7) or anti-cZP1 N-terminal fragment polyclonal^57^ (1:3,000). Secondary antibodies were horseradish peroxidase-conjugated goat anti-mouse (1:10,000) (Life Technologies/Thermo Fisher Scientific) or horseradish peroxidase-conjugated horse anti-mouse (1:10,000) (Cell Signalling Technology). Chemiluminescence detection was performed with Western Lightning ECL Plus (Perkin Elmer).

For cell lysate analysis, cells transfected for 72 h were washed twice with 1 mL cold phosphate-buffered saline (PBS). 200 µL lysis buffer containing 50 mM Na- HEPES pH 7.8, 150 mM NaCl, 0.1% (v/v) SDS, 0.5% (v/v) sodium deoxycholate, 1% (v/v) Triton X-100 and protease inhibitors (cOmplete mini EDTA-free protease inhibitor cocktail; Roche) were added to one well and the plate incubated overnight at 4°C. Lysate samples were recovered by scraping cells in the presence of SDS-PAGE loading buffer. 40 µL cell lysate was boiled and separated by SDS-PAGE for immunoblot analysis.

Deglycosylation experiments in reducing conditions were performed with PNGase F (New England Biolabs), according to the manufacturer’s instructions; for analysing deglycosylation products in non-reducing conditions, DTT-containing Glycoprotein Denaturing Buffer was replaced with 10 mM Na-HEPES pH 7.5, 1% (v/v) SDS.

### Preparation of soluble ZP glycoproteins from proteolytically degraded chicken egg coat

Egg coats were isolated from the F1 oocytes of laying white leghorn hens, and sperm as well as sperm protease prepared from ejaculated semen as described previously^44^. 100 mg isolated egg coat (wet weight) were suspended in 1 mL PBS by ultra-sonication^31^ and then mixed with the sperm protease preparation. The reaction mixture was kept at 39°C for 20 h and centrifuged at 16,000 × g for 5 min. The supernatant was 0.45-μm filtered and injected into a HiPrep 16/60 Sephacryl S-300 HR column (GE Healthcare). SEC elution profiles were monitored by measuring absorbance at λ=280 nm, and both the ∼35 kDa and ∼21 kDa N-terminal fragments of cZP1 were analysed by mass spectrometry following in-gel chymotryptic peptide digestion as previously described^60^. Animal experiments were performed in accordance to the local ethics committee’s approval (Graduate School of Bioagricultural Sciences, Nagoya University, approval number 2016030218).

### Mass spectrometry

For LC-MALDI-TOF MS/MS analysis, N-terminal fragments of native cZP1 were boiled in SDS in the presence of 0.75% (v/v) β-mercaptoethanol and treated with PNGase. Both deglycosylated and glycosylated samples were reduced, alkylated and digested with chymotrypsin in the presence of 0.01% (w/v) ProteaseMAX™ Surfactant (Promega). A part of these peptide preparations was subjected to LC-MALDI-TOF MS/MS analysis. Chymotryptic glycopeptides from the sample without PNGase treatment were captured using RCA-1 lectin, and the resulting complexes concentrated with 30 kDa cut-off centrifugal filtration devices. Bound glycopeptides were released using 0.1% (v/v) trifluoroacetic acid and analysed by MALDI-TOF/MS.

Samples were injected into a DiNa Nano LC system equipped with a DiNa MaP autospotter (KYA technologies) and eluted using a linear acetonitrile gradient. MS spectra were obtained using a MALDI-TOF/TOF 5800 Proteomic Analyser mass spectrometer (Applied Biosystems) and analysed using Mascot^61^.

### Protein purification

72 h after transfection, the conditioned media from mammalian cells was harvested, 0.22 μm-filtered (Sarstedt) and adjusted to 20 mM Na-HEPES pH 7.8, 500 mM NaCl, 5-10 mM imidazole (IMAC buffer). 10 mL pre-equilibrated nickel agarose slurry (Ni-NTA; QIAGEN) was added per litre of medium and incubated for either 1 h at room temperature (RT) or overnight at 4°C. After washing the beads with 100 column volumes IMAC buffer, proteins were batch-eluted with 5 column volumes elution buffer (20 mM Na-HEPES pH 7.8, 150 mM NaCl, 500 mM imidazole). HEK293S-expressed cZP1-N1 was deglycosylated with Endo H (1:10 mass ratio) for 1 h at 37°C in 120 mM Na/K phosphate pH 6.0 prior to elution.

Proteins were concentrated with appropriate centrifugal filtration devices (Amicon) and further purified by SEC at 4°C using an ÄKTA_FPLC_ chromatography system (GE Healthcare). For crystallographic studies, cZP1-N1 was injected into a HiLoad 26/600 Superdex 200 pg column (GE Healthcare) pre-equilibrated with 20 mM Na-HEPES pH 8.0, 500 mM NaCl. Protein-containing fractions were pooled and concentrated to 5 mg/ml (Endo H-deglycosylated cZP1-N1) or 3.5 mg/ml (glycosylated cZP1-N1), respectively; prior to crystallization setup, samples were dialyzed against a buffer consisting of 20 mM Na-HEPES pH 8.0, 200 mM NaCl. For analytical SEC and SEC-MALS experiments, 20 mM Na-HEPES pH 7.8, 150 mM NaCl was used as running buffer.

### Sequence analysis

To create a non-redundant database of ZP1-N1 sequences, a redundant collection of 510 entries was first created by combining the results of parallel BLAST^62^ searches performed using as queries a selection of ZP1 homologs from HomoloGene^63^ entry 33483. After pruning of sequences identical to or contained within others, 128 entries with 5 Cys were processed with CD-HIT^64^, using a 0.8 sequence identity threshold. Following manual restore of the mZP1-N1 sequence (belonging to the same CD-HIT cluster as hZP1-N1) and deletion of 2 sequences whose full-length counterparts did not contain a trefoil domain^65,66^, this resulted in a non-redundant database of 28 sequences from amniotes which were aligned with MAFFT^67^ as implemented in SnapGene (GSL Biotech).

### SEC-MALS analysis

Measurements were performed using an Ettan LC high-performance liquid chromatography system with a UV-900 detector (Amersham Pharmacia Biotech; λ=280 nM), coupled with a miniDawn Treos MALS detector (Wyatt Technology; λ=658 nm) and an Optilab T-rEX dRI detector (Wyatt Technology; λ=660 nm). Separation was performed using a Superdex 75 Increase 10/300 GL column (GE Healthcare), using a flow rate of 0.4 ml/min and a mobile phase consisting of 10 mM Na-HEPES pH 7.8, 150 mM NaCl. Data processing and weight-averaged molecular mass calculation were performed using the ASTRA 7.1.3 software (Wyatt Technology).

### Protein crystallization

Hanging drop vapor diffusion experiments were set up by hand at 20°C, by mixing protein and reservoir solutions at a 1:1 ratio and equilibrating against 1 mL reservoir solution. Endo H-treated cZP1-N1 N121Q from HEK293S cells and untreated cZP1-N1 from HEK293T cells crystallized in 0.6 M LiCl, 0.1 M tri-sodium citrate pH 5.5 and 29% (v/v) MPD, 0.1 M MES pH 6.5, respectively. For heavy-atom derivatization, crystals of the Endo H-treated protein were soaked in reservoir solutions supplemented with 5 mM KAu(CN)_2_ or 1 mM Zn(OAc)_2_ for 20 h before cryocooling.

### X-ray data collection

All datasets were collected at 100 K from single crystals cryoprotected by supplementing the mother liquor with MPD to a final concentration of 35% (v/v) and subsequently flash-frozen in liquid nitrogen.

The native dataset for glycosylated cZP1-N1 was collected at European Synchrotron Radiation Facility (ESRF, Grenoble) beamline ID23-1^68^ (λ=0.9791 Å), using a PILATUS 6M detector (DECTRIS). Datasets for the Endo H-treated protein were collected at beamline BL14.1 of BESSY II (Helmholtz-Zentrum, Berlin) using a MarMosaic 225 CCD detector (Mar) (native data, λ=1.7710 Å) or at beamline I03 of Diamond Light Source (DLS, England) using a a PILATUS 6M-F detector (DECTRIS) (gold derivative data, λ=1.03970 Å; zinc derivative data; λ=1.2825 Å). Data collection statistics generated using phenix.merging_statistics^69^ are reported in Table 1.

### Data processing and structure determination

X-ray diffraction datasets were processed using XDS^70^.

The structure of the gold derivative of Endo H-treated cZP1-N1 was solved at 2.75 Å resolution by single-wavelength anomalous dispersion (SAD) phasing with PHENIX AutoSol^71^, which located 6 sites (FOM 0.35, BAYES-CC 53.5). The resulting initial model, which included both moieties of a ZP1 cross-link for a total of 155 residues (R_work_ 42%, R_free_ 44%, model-map correlation 62%), was manually rebuilt in Coot^72^ and refined against the 2.3 Å resolution native dataset using phenix.refine^73^; protein geometry was validated with MolProbity^74^ and carbohydrate structure validation was carried out using Privateer^75^. The structure of the same protein was also independently solved by Zn-SAD phasing at 2.7 Å resolution (FOM 0.31, BAYES-CC 45.1), yielding an initial model of 163 residues (R_work_ 35%, R_free_ 0.41%, model-map correlation 73%) that was manually completed and led to a refined set of coordinates that includes 9 Zn^2+^ ions. The structure of glycosylated cZP1-N1 was determined by molecular replacement (MR) with Phaser^76^, using as search model the structure of a monomer of Endo-H treated cZP1-N1.

The MolProbity Ramachandran plots of the structures described in this study show the following percentages of favoured, allowed and outlier residues: 98.6, 1.4, 0.0 (Endo H-treated cZP1-N1); 99.5, 0.5, 0.0 (Endo H-treated cZP1-N1 with Zn^2+^); 100.0, 0.0, 0.0 (glycosylated cZP1-N1). Refinement and validation statistics generated using phenix.table_one^69^ are reported in Table 1.

### Structure analysis

Structural alignments were performed using UCSF Chimera^77^; protein-protein interfaces were analysed manually as well as using FoldX^78^, PDBsum^79^, PIC^80^ and PISA^81^. Electrostatic surface potential calculations were performed with PDB2PQR^82^ and APBS^83^, via the APBS Tools plugin of PyMOL (Schrödinger, LLC). Figures were created with PyMOL.

### Molecular dynamics simulations

Simulations were performed using GROMACS version 2016^84^ and the CHARMM36 force field^85,86^, based on the crystallographic coordinates of the Zn^2+^-bound cZP1-N1 homodimer. Simulation conditions were set up essentially as described^29^, using a 8 nm dodecahedric box and 0.1 M Na+/Cl-ion concentrations. 3 independent 100 ns replicas were performed; before data generation, 20-ns position-restrained simulations were performed to relax the positions of Zn^2+^-proximal water molecules.

### Pull-down analysis of protein-protein interaction

For pull-down experiments of 6His-tagged hZP4-N1 and cZP4-N1 co-transfected with Myc-tagged counterparts, 2 mL conditioned media were harvested, equilibrated with 20 mM Na-HEPES pH 7.8, 100 mM NaCl, 10 mM imidazole and incubated with 50 μL Ni-NTA beads for 1 h at RT. Beads were collected by centrifugation at 100 × g and washed 4 times with 500 μL binding buffer followed by centrifugation at 100 × g for 5 min; bound proteins were eluted with 100 μL 20 mM Na-HEPES pH 7.8, 150 mM NaCl, 500 mM imidazole. 100 μL flow-through and 5 μL elution fractions were treated with PNGase F using standard procedures and analysed by immunoblotting with anti-5His and anti-Myc.

## Data availability

Atomic coordinates and structure factors have been deposited with the Protein Data Bank (PDB) under accession codes 6GF6 (Endo H-treated cZP1-N1 homodimer, high resolution native), 6GF7 (Endo H-treated cZP1-N1 homodimer, Zn^2+^ derivative) and 6GF8 (glycosylated cZP1-N1 homodimer). A reporting summary for this article is available as a Supplementary Information file.

## Acknowledgments

This work was supported by Karolinska Institutet; the Center for Biosciences; the Center for Innovative Medicine; Swedish Research Council grants 2012-5093 and 2016-03999; the Göran Gustafsson Foundation for Research in Natural Sciences and Medicine; the Sven and Ebba-Christina Hagberg foundation; an EMBO Young Investigator award; and the European Research Council under the European Union’s Seventh Framework Programme (FP7/2007-2013)/ERC grant agreement 260759 (L.J.); JSPS KAKENHI grants 22112510 and 17380200 (T.M.). We are grateful to ESRF, DIAMOND and HZB synchrotrons for beamtime allocation and assistance with data collection; R. Aricescu (MRC Laboratory of Molecular Biology) for HEK293T cells; D. Leahy (Johns Hopkins University School of Medicine) for *E. coli* expression vector pProEX HT-endoglycosidase H. We also thank Dirk Fahrenkamp for help with SEC-MALS analysis and other members of the Jovine laboratory for discussion and comments.

## Author contributions

K.N. expressed, purified and crystallized proteins; determined and refined structures together with L.J. and performed part of the biochemical assays. E.D. carried out all other biochemical and biophysical studies. S.N. and T.M. analysed native cZP1 biochemically and by mass spectrometry. L.H. and S.N. helped with mammalian expression. A.V. performed and analysed MD simulations. L.J. directed the study and wrote the manuscript with E.D. and K.N.

## Additional information

### Competing interests

the authors declare no competing financial interests.

---

**Supplementary Figure 1.**
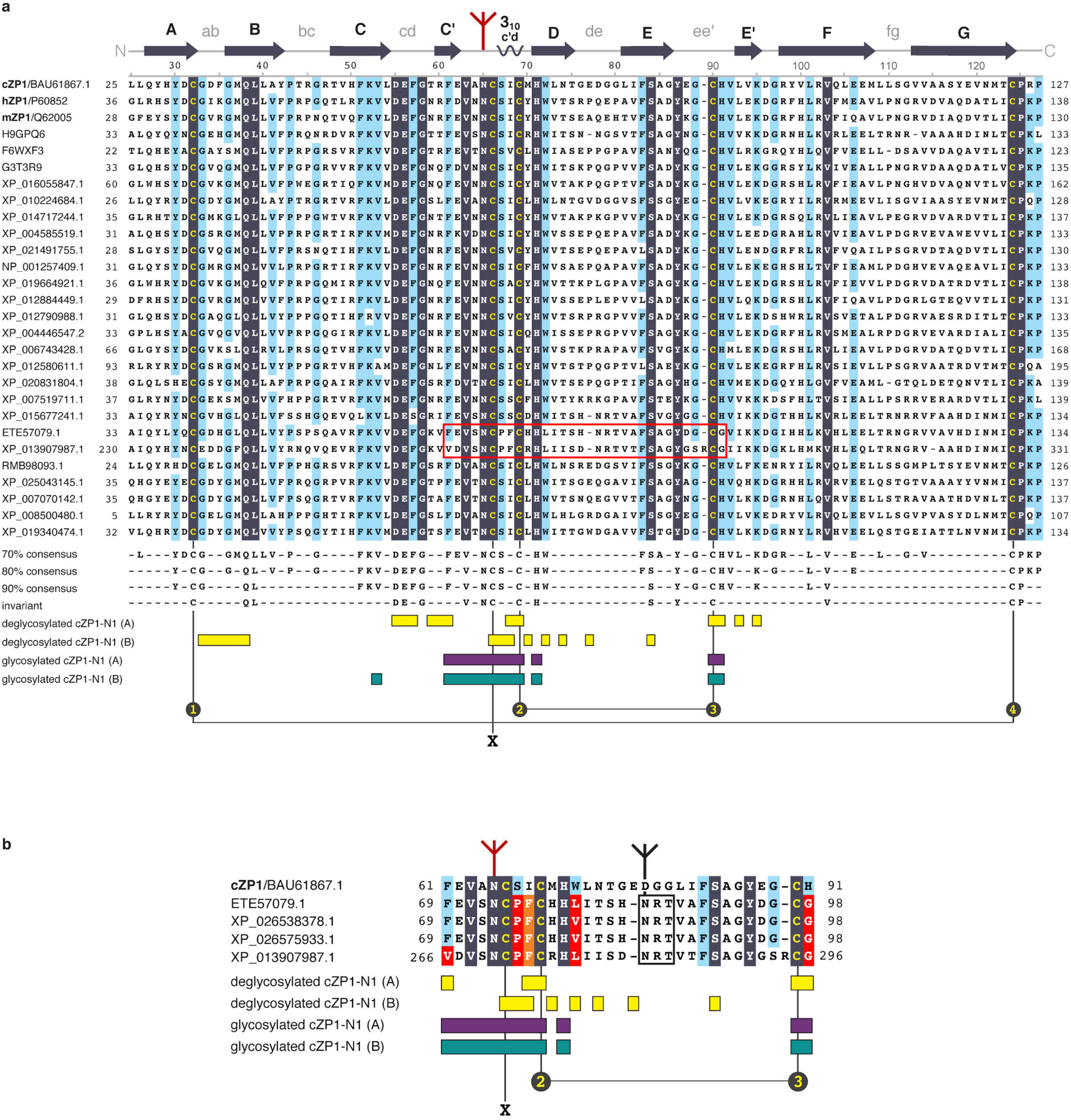
Sequence alignment of representative ZP1-N1 sequences. **a**, Alignment of a 80% identity non-redundant ZP1-N1 sequence database, with invariant residues shaded in dark blue and amino acids above the 80% identity cut-off shaded in cyan. Cys residues are highlighted in yellow. Sequences are indicated by accession numbers and associated amino acid boundaries. Above the alignment, the residue numbering and secondary structure diagram of cZP1 are reported. β-strands are indicated by arrows, whereas the 3_10_ helix in the c’d loop (only found in chain A of deglycosylated cZP1-N1 as well as chain B of the glycosylated protein) is depicted as a squiggle. An inverted dark red tripod symbolizes the glycan attached to the highly conserved N-glycosylation site of ZP1-N1. Below the alignment, residues matching different identity thresholds within the 100%-70% range are shown, and positions corresponding to amino acids involved in the different cZP1-N1 homodimer interfaces are indicated by bars, coloured according to Fig. 5. The canonical ZP-N intramolecular disulphide pattern is also shown, and the ZP1-specific invariant Cys forming the intermolecular cross-link is indicated by the X symbol. **b**, Expansion of the alignment region contained within the red rectangle in panel a to include all members from two clusters consisting of sequences from the suborder Serpentes. The corresponding region of the cZP1 sequence is also reported for comparison, and elements are represented as in panel a. Notable variations in asymmetric or symmetric homodimer interface residues are highlighted using red or orange shading, respectively. A black rectangle marks the sequon that replaces the N-glycosylation site corresponding to cZP1 N65 (inverted dark red tripod) with one located in the de loop (inverted black tripod).

**Supplementary Figure 2.**
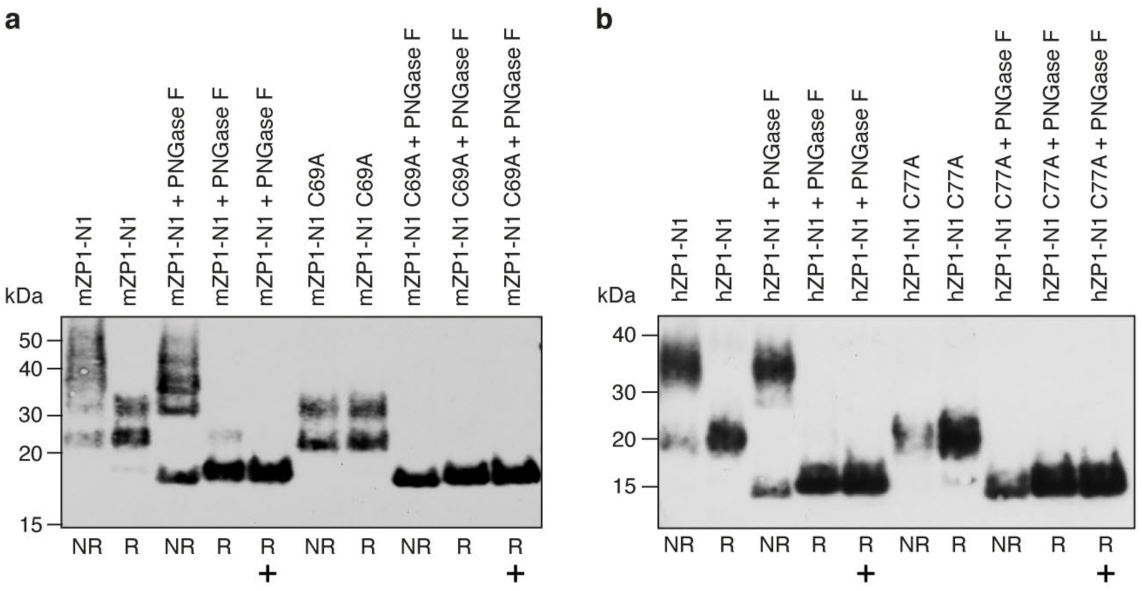
Anti-5His immunoblot analysis of purified mZP1-N1 (**a**) and hZP1-N1 (**b**) proteins before and after deglycosylation with PNGase F. The experiment was performed using the material shown in Fig. 6e, g, and samples digested using the manufacturer’s Glycoprotein Denaturing Buffer, which contains DTT, are indicated by a plus sign. Note that PNGase F is unable to efficiently deglycosylate the homodimeric form of the mammalian proteins in non-reducing conditions; this suggests that, like cZP1 N65, N76 of hZP1 and N68 of mZP1 also lie in close proximity to the cross-link interface. Source data are provided as a Source Data file.

**Supplementary Figure 3.**
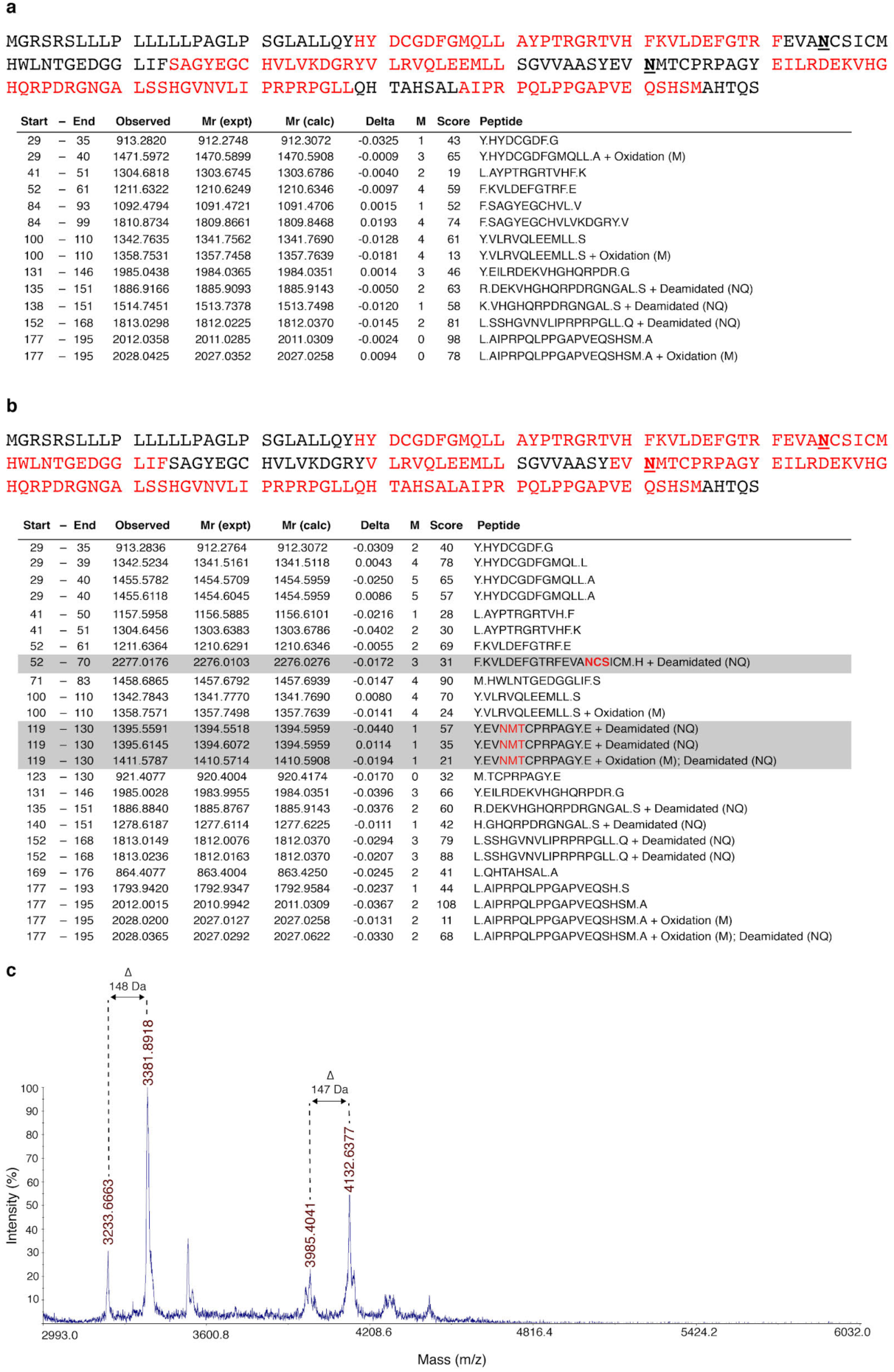
Mass spectrometry analysis of native chicken ZP1 N-terminal fragment. **a-b**, Soluble material released by sperm protease treatment of native chicken egg coats was reduced, digested with chymotrypsin and subjected to LC-MALDI-TOF MS/MS analysis. Sequence coverage by MS analysis is indicated in red and N-glycosylation sites are underlined. Whereas cZP1 N-terminal fragment peptides including N65 or N121 are missing in the fully glycosylated sample (a), the material treated with PNGase F shows peptides containing N65 or N121 (b, grey highlight). **c**, Chymotryptic glycopeptides were enriched using RCA-1 lectin and analysed by MALDI-TOF MS analysis. Four major peaks above 3000 m/z were detected: the first two (3233.6663 and 3381.3193 m/z) are consistent with a glycopeptide containing the N-glycan linked to N121 of the peptide E119-Y130 (Mr, 1396), whereas the 3985.4041 and 4132.6377 m/z peaks are consistent with a peptide carrying a carbohydrate linked to N65 of the peptide K52-M70 (Mr, 2277). The differences between these pairs of peaks suggest that they differ by a single fucose residue (mass 146 Da).

**Supplementary Figure 4.**
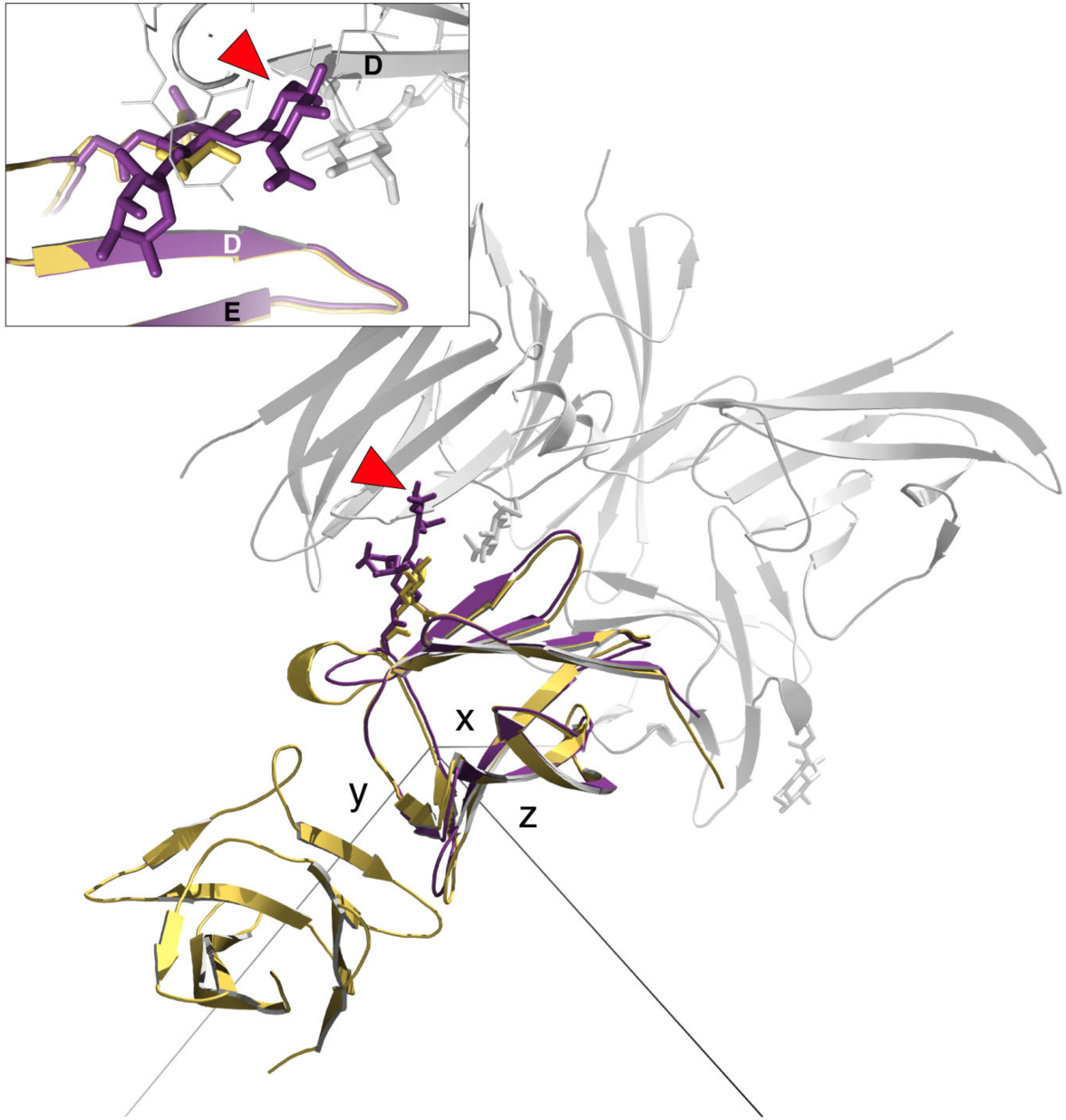
Superimposition of glycosylated and Endo H-deglycosylated cZP1-N1 (violet purple and yellow, respectively). Presence of an α1- 6-linked fucose (red arrowhead) would not be compatible with the packing of the tetragonal crystal form because it would clash with β-strand D of a symmetry-related molecule (grey).

**Supplementary Figure 5.**
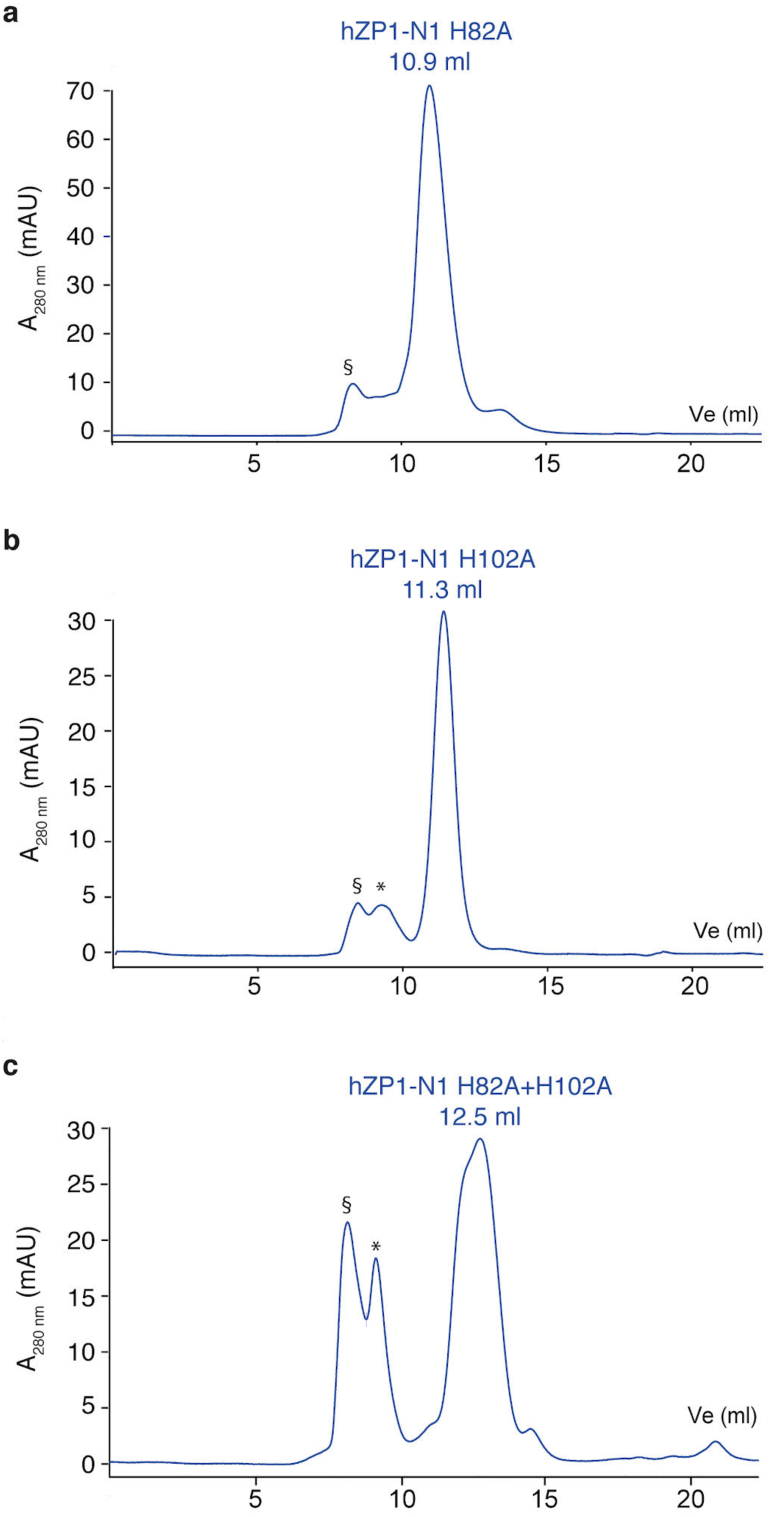
SEC Analysis of hZP1-N1 mutants. **a-c**, SEC profiles of hZP1-N1 mutants H82A (a), H102A (b) and hZP1-N1 H82A+H102A (c) using a Superdex 75 Increase 10/300 GL column. A Coomassie-stained SDS-PAGE analysis of the corresponding peaks fractions is shown in Fig. 8h. Elution volumes are reported; void volume and contaminant peaks are indicated by § and *, respectively.

**Supplementary Table 1.** Marsupial and eutherian ZP properties

|  | ZP composition |  |  |  | ZP thickness |
| --- | --- | --- | --- | --- | --- |
|  | ZP1 | ZP2 | ZP3 | ZP4 |  |
| Gray short tail opossum<sup>87</sup><br>(*Monodelphis domestica*) | + | + | pseudogene ZP3a<br>(ZP3b, c) | pseudogene | 1.6-2.0<sup>88</sup> $\mu\text{m}$ 0.5 $\mu\text{m}$ -1 $\mu\text{m}$<sup>89</sup><br>(no more than 4 $\mu\text{m}$<sup>90-92</sup>) |
| Brushtail possum<br>(*Trichosurus vulpecula*) | + | + | + | + | 8 $\mu\text{m}$<sup>93</sup> |
| Tammar Wallaby<sup>87</sup><br>(*Macropus eugenii*) | + | + | (ZP3a, b, c) | + | 4-8.5 $\mu\text{m}$<sup>93</sup> |
| Mouse<br>(*Mus musculus*) | + | + | + | *pseudogene* | 5-7 $\mu\text{m}$<sup>94</sup> |
| Hamster<br>(*Mesocricetus auratus*) | + | + | + | + | 10-16 $\mu\text{m}$<sup>95</sup> |
| Human<br>(*Homo sapiens*) | + | + | + | + | 15-20 $\mu\text{m}$<sup>21</sup> |
| Rabbit<br>(*Oryctolagus cuniculus*) | + | + | + | + | 16 $\mu\text{m}$<sup>96</sup> |
| Cat<br>(*Felis catus*) | + | + | + | + | 15 $\mu\text{m}$<sup>97</sup> |
| Dog<br>(*Canis lupus*) | - | + | + | + | 10-15 $\mu\text{m}$<sup>98</sup> |
| Pig<br>(*Sus scrofa*) | - | + | + | + | 16 $\mu\text{m}$<sup>99</sup> |
| Cow<br>(*Bos taurus*) | - | + | + | + | 27 $\mu\text{m}$<sup>99</sup> |
| Horse<br>(*Equus caballus*) | + | + | + | + | 8 $\mu\text{m}$<sup>97</sup> |

